# Demographic history inferred from an inversion-rich spruce bark beetle genome

**DOI:** 10.1101/2025.07.17.665273

**Authors:** Piotr Zieliński, Julia Morales-García, Martin Schebeck, Mihai Leonard Duduman, Krystyna Nadachowska-Brzyska

## Abstract

The demographic history of species inferred from whole-genome data provides quantitative insights into key biological parameters such as population size changes and divergence times. Reliable estimates often require data that have not been affected by selection. Extensive research, however, indicates that many species harbour multiple polymorphic chromosomal inversions, which often evolve under different selective pressures. Consequently, inversions can influence genome-wide patterns of variation and subsequent evolutionary inferences. In this study, we used genome-wide data from over 300 spruce bark beetle (*Ips typographus*) individuals from 23 populations across Europe to reconstruct their demographic history and to investigate the impact of a complex polymorphic inversion landscape (covering approximately 28% of the beetle genome) on demographic inference. We used two complementary methods, Pairwise Sequential Markovian Coalescent (PSMC) and Site Frequency Spectrum (SFS)-based modelling, and revealed a Late Pleistocene divergence (∼79 kya) between populations from the southern and northern parts of the species’ European range, and a long-term effective population size of ∼250,000. The southern group underwent significant population expansion after this divergence event, whereas the northern group expanded during the Holocene (∼7 kya). Recent population size estimates suggest that the southern group is twice as large as the northern group. Neglecting the presence of chromosomal inversions did not significantly affect the model selection procedure and resulted in relatively small biases in the estimated demographic parameters. This study provides information on the historical population dynamics of the spruce bark beetle and improves our understanding of the influence of a complex genomic architecture on the inference of evolutionary history.

## Introduction

Demographic history is reflected in the patterns of genetic variation observed within and among populations. Therefore, demographic models can complement paleontological data by providing information on biologically relevant parameters, such as effective population size, time of divergence and migration rate. For decades, demographic analysis of populations has relied on a small fraction of the genome, e.g. genetic markers such as mtDNA or a limited set of nuclear loci (Hey, 1997; Nielsen, 2000). The advent of high-throughput sequencing has revolutionised approaches to infer the demographic history of numerous species (Gutenkunst et al., 2009; Leuenberger & Wegmann, 2010; Terhorst et al., 2017). For example, new methods have been developed to use genomes of single individuals to infer historical demographic events (Gronau et al., 2011; Li & Durbin, 2011). Improvements have also been made to classical approaches to incorporate large genome-wide datasets (Excoffier et al., 2013; Schiffels & Durbin, 2014; Speidel et al., 2019). While these developments provide greater power and resolution to estimate population parameters, they also pose new challenges (Marchi et al., 2021). As they require, for example, consideration of genome-wide selective pressures. Excluding loci that could potentially evolve under selection, such as coding regions, is common practice in demographic inference (Johri et al., 2021). There is also a growing body of research on the effects of genome-wide background selection (Charlesworth, 2013; Ewing & Jensen, 2016; Johri et al., 2021). However, the influence of large polymorphic rearrangements on demographic inference remains unclear.

This is important because many genomic rearrangements evolve under different selective pressures (Berdan et al., 2023; Faria, Johannesson, et al., 2019) and often cover large parts of the genome (Barb et al., 2014; Faria, Chaube, et al., 2019; Harringmeyer & Hoekstra, 2022). Chromosomal inversions, which occur when a segment of a chromosome breaks off, changes orientation and reconnects in the same place, can have a particularly complex effect on genome-wide patterns of variation. When an inversion occurs, it creates a polymorphism with two haplotypes (inverted and collinear) and effectively suppresses recombination in heterozygous individuals (Wellenreuther & Bernatchez, 2018). As a result, the two inversion haplotypes evolve largely independently, accumulating new mutations separately, and alleles present on the same inversion haplotype are maintained in strong linkage disequilibrium (Berdan et al., 2023; Faria, Johannesson, et al., 2019; Kirkpatrick, 2010; Mykhailenko et al., 2024). The segregation into two inversion haplotype ‘populations’ ultimately reduces the effective population size of each haplotype and creates a hidden population structure present only in the affected part of the genome (Berdan et al., 2021). As a result, this part of the genome may be characterised by patterns of variation consistent with a different demographic history than collinear parts of the genome (Novo et al., 2023; Poikela et al., 2024).

Importantly, the effect of inversions on demographic inference depends on several factors and requires empirical testing to quantify the magnitude of the expected biases and to identify the scenarios that may lead to false results, e.g. incorrect model choice or parameters that have been over- or under-estimated. Key factors include the proportion of the genome affected by rearrangements, the frequency of inversions, the nature of selective pressures acting on/within inversions, the age of inversions, and their distribution across the species range (Berdan et al., 2023; Berdan et al., 2021; Faria, Johannesson, et al., 2019; Seich al Basatena et al., 2013; Wellenreuther & Bernatchez, 2018). We expect that inferences based on genome-wide data should not be biased if the size of the inversion(s) is relatively small (Connallon & Olito, 2022). If one of the inversion haplotypes is rare, the effect of reduced recombination, determined by the frequency of heterozygotes, will also be limited (since most individuals have two copies of the common inversion haplotype). However, if the inversion haplotypes have comparable frequencies, for example if they evolve under balancing selection and have been present in the population for a considerable period of time, this will result in an excess of divergent haplotypes with high-frequency alleles, which can falsely suggest population contraction (Corbett-Detig & Hartl, 2012; Navarro et al., 2000). If positive selection has been acting within a particular inversion haplotype, the signature of a selective sweep will spread across the entire inversion haplotype, leading to a greater reduction in variation compared to a selective sweep in regions where recombination has not been restricted (Begun & Aquadro, 1992; Cutter & Payseur, 2013; Kennington & Hoffmann, 2013). In particular, genetic hitch-hiking of neutral alleles associated with a beneficial mutation undergoing a selective sweep, or selective removal of deleterious mutations by background selection, will have a greater effect on patterns of polymorphism in genomic regions with restricted recombination (Charlesworth et al., 1993; Kaplan et al., 1989), this in turn may lead to lower estimates of effective population size. Furthermore, the effects of older inversions may be more pronounced than those of more recent origin, simply because there would have been more time for divergence between inversion haplotypes to build up and/or more time for selective pressure to influence the observed variation (Kapun & Flatt, 2019). In addition, the geographic distribution of inversion haplotypes may influence their impact on demographic inference. For example, polymorphic inversions that evolve under balancing selection and are present across the species range may inflate estimates of gene flow between populations, when in fact they are ancestral polymorphisms that have segregated within the species for longer than expected under neutrality (Wellenreuther & Bernatchez, 2018). Conversely, inferences based on inversion haplotypes that evolve under divergent selection (alternative haplotypes are favoured in different ecological niches) may underestimate gene flow between these environments (Barth et al., 2019). Finally, differences in the sensitivity of demographic methods to the effects of inversions may be expected if the methods use different types of data or summarise genomic data differently e.g. single whole genome sequences vs. site frequency spectra from multiple individuals (Beichman et al., 2017).

In this study, a Pairwise Sequential Markovian Coalescent (PSMC) and Site Frequency Spectrum (SFS) based approach was used to infer the demographic history of the Eurasian spruce bark beetle, *Ips typographus*. This species is one of the most destructive forest pests that exhibits eruptive population outbreaks under suitable environmental conditions, causing large-scale mortality of spruce-dominated forests. The economic importance of spruce has led to extensive research on the ecology, biology and impacts of the spruce bark beetle on forests and society, but genetic studies have until recently been limited to the analysis of a few genetic markers (Bertheau et al., 2013; Mayer et al., 2015; Papek et al., 2024; Stauffer et al., 1999). A recent genome re-sequencing study revealed that the spruce bark beetle genome harbours one of the most complex polymorphic inversion landscapes described to date (Mykhailenko et al., 2024). Twenty-seven large polymorphic inversions, covering approximately 28% of the species’ genome, make the spruce bark beetle genomic data ideal for exploring the effects of inversions on demographic inference. Here we aim to: a) infer the unknown demographic history of the spruce bark beetle by considering regions affected by inversions, b) assess whether neglecting the presence of inversions introduces biases in parameter estimates and the demographic models chosen, and c) test whether different inference methods are equally affected by the presence of inversions.

## Materials and methods

### Sampling, sequencing and data preparation

We build on the dataset presented in Mykhailenko et al. (2024), where 13-14 *I. typographus* individuals were sampled from 18 populations distributed across the species’ European range. In addition, we sampled 72 individuals from five populations in the Carpathians, the Alps, and the Apennines. In total, the dataset consisted of 23 populations and 312 individuals (Figure 1, Table S1). As in our previous study, DNA from the new samples was extracted from whole beetles using the Wizard Genomic DNA Purification Kit (Promega) and subjected to whole-genome re-sequencing (2×150 bp paired-end NovaSeq 6000 Illumina sequencing). Raw sequencing data were processed as described in Mykhailenko et al. (2024). Briefly, low quality reads and adaptors were trimmed using Trimmomatic 0.39 (Bolger et al., 2014), data were mapped to the *I. typographus* genome (Powell et al., 2021) using Bowtie2 version 2.4.2 (Langmead & Salzberg, 2012), and duplicate reads were removed using PicardMarkDuplicates version 2.24.1 (Picard). Mappings were recalibrated using GATK version 4.1.9.0 BaseRecalibrator and ApplyBQSR (Depristo et al., 2011; McKenna et al., 2010). Variant calling, genotyping, recalibration and filtering were performed using GATK HaplotypeCaller, GenotypeGVCFs, VariantRecalibrator, and VariantFiltration, respectively. Five sites around indels were filtered out using BCFtools version 1.11 (Danecek et al., 2021). To remove low quality variants and sites that may come from duplicated regions, we filtered out variants with a mapping quality of less than 30, a total depth (calculated for all individuals) greater than the mean + one standard deviation, variants with excessive heterozygosity, and all sites within repetitive regions as identified by Powell et al. (2021). Individual genotypes with a coverage less than 8 and genotype quality less than 20 were treated as missing. Finally, we retained biallelic SNPs with less than 50% of missing genotypes. Similar to Mykhailenko et al. (2024), we focused on contigs larger than 1Mb (36/276, covering 78% of the genome assembly). However, we also excluded IpsContig33, which has a high similarity to bark beetle mtDNA in part of its length, which may be due to some assembly errors, and IpsContig9 which is most likely involved in sex determination as it is hemiploid in males (Mykhailenko et al., 2024). In addition, we also excluded several short contigs with a low number of SNPs (number of SNPs < 10k, π_S_ < 0.007, π_S_ < mean π_S_ / 4), treating them as non-informative compared to the rest of the genome. Thus, our final dataset consisted of variation data calculated over 25 contigs spanning 163 Mb, representing 69% of the reference genome.

**Figure 1.**
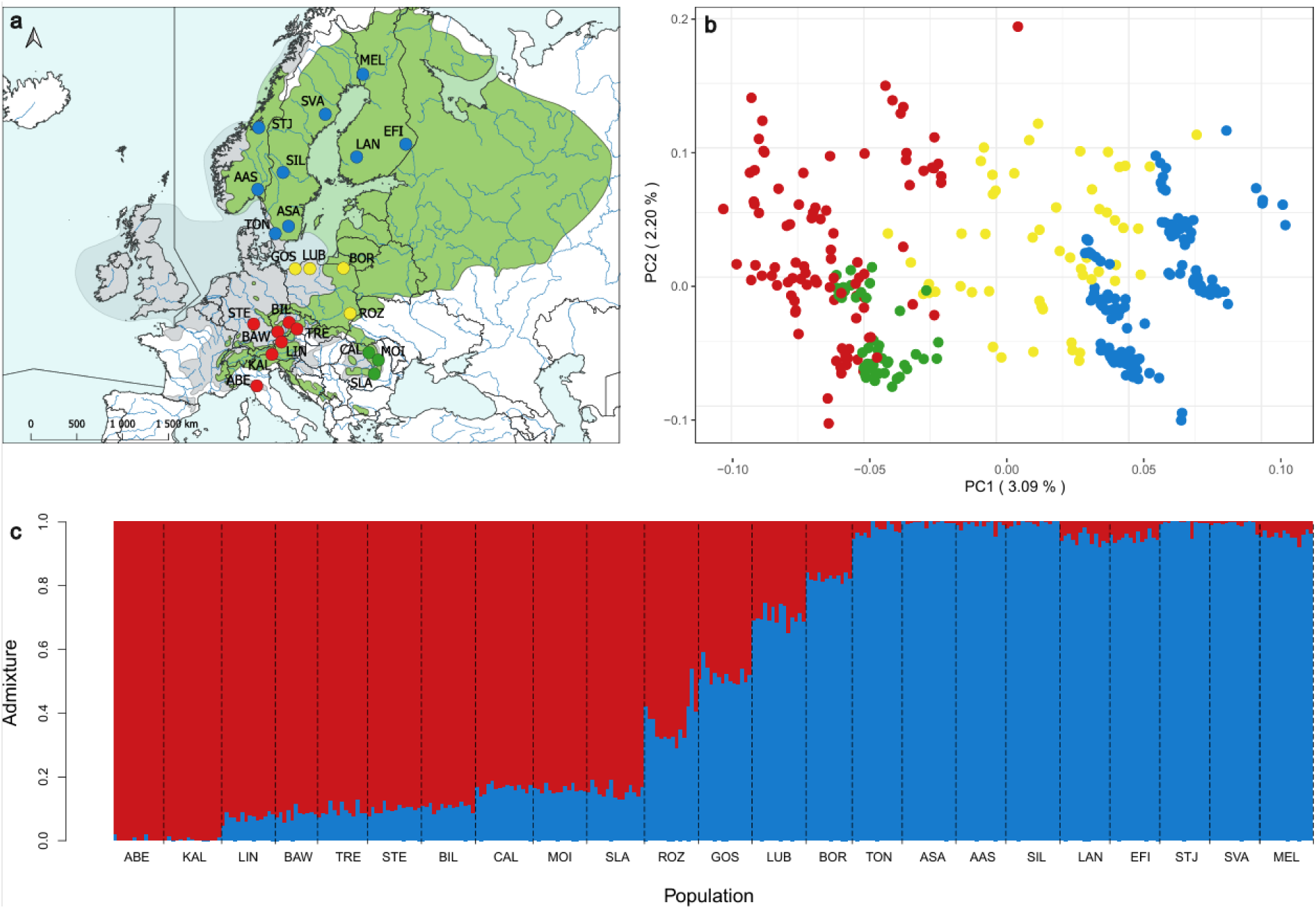
Sampling sites and intraspecific genetic structuring. a) Sampling locations. Blue - populations assigned to the northern group, red - southern group, yellow - Polish populations, green - Carpathian populations. b) PCA based on genome-wide variation data. Colour scheme as in a. c) Individual admixture proportions for K=2. Population abbreviations as assigned in Table S1.

### Genetic structure and genome-wide variation

To determine the genome-wide genetic structure of the species, including newly sampled populations, we performed a PCA on the whole-genome and contig by contig using PLINK version 1.90b6.18 (Purcell et al. 2007). To determine the number of genetic clusters (K) and admixture proportions, we performed NGSAdmix analysis (Skotte et al., 2013) using LD-truncated data (K=1-5; 10 replicates per K value, MAF=0.05 and 10,000 iterations), followed by likelihood and CLUMPAK analysis (Kopelman et al., 2015) to select the most likely number of clusters. Finally, we calculated pairwise *F_ST_* values to test for genetic differentiation between populations. To test whether inversions are in Hardy-Weinberg (HW) equilibrium, we used Genepop version 4.1.2 (Rousset, 2008), controlling for type I error using the false discovery rate (FDR) procedure (Benjamini & Hochberg, 1995). To assess if and how genetic variation is affected by the presence of polymorphic inversions, we calculated several standard population genetic measures, i.e. nucleotide diversity (π), Tajima’s D, linkage disequilibrium (LD), *F_ST_* and *d_XY_*. Nucleotide diversity, Tajima’s D and *d_XY_* were calculated using ANGSD version 0.935-44-g02a07fc (Korneliussen et al., 2014). *F_ST_* values were calculated in VCFtools version 0.1.16 (Danecek et al., 2011) while LD using PLINK version 1.9 (Chang et al., 2015). Statistics were calculated in 100 kb windows. The results were then compared across the datasets (see next section). The statistical significance of the differences was tested with 1,000 permutations.

### Pairwise Sequential Markovian Coalescent (PSMC) analysis

Effective population size changes over time were inferred using PSMC (Li & Durbin, 2011). PSMC relies on the distribution of polymorphic sites across a single genome, and its performance is sensitive to differences in genome coverage (Nadachowska-Brzyska et al., 2016). Therefore, we decided to standardise coverage across samples to a level of 25x (average coverage per base) and only analysed individuals that met this criterion (129 individuals in total). We used Qualimap version 2.2.2 (Okonechnikov et al., 2016) to measure coverage per site, and Picard DownsampleSam to subsample all individuals to the same coverage. Consensus sequences per individual were obtained with Samtools mpileup version 1.11, BCFtools call, and vcfutils vcf2fq (Danecek et al., 2021). Sites with coverage lower than 8 (mean/3) and higher than 50 (2 x mean) were masked for downstream analysis. To assess the influence of inversions on PSMC inference, we performed analyses on four datasets: 1) whole-genome (including inversions), 2) no-inversions (only the collinear part of the genome; 72% of the analysed part of the reference genome), 3) inversions-only (genome sequences excluding collinear parts of the genome; 28% of the analysed part of the genome), 4) to assess if the size of the inversions-only dataset itself could affect obtained Ne trajectories we run PSMC also on the collinear part of the genome trimmed to the size of the inversion-only dataset (no-inversions-small dataset), keeping the same fragment sizes as for regions affected by inversions. The PSMC analysis settings (-N25 -t10 -r5 -p “4+25*2+4+6”) were chosen manually following the suggestions by Li and Durbin (https://github.com/lh3/psmc) and were kept the same for all individuals and datasets. A generation time of 1 year was used for all populations, despite current differences in species distribution. Currently, northern populations typically have one generation per year, while southern populations usually have two, under favourable conditions up to three generations per year (Annila, 1969; Heliövaara & Peltonen, 1999; Jönsson et al., 2009; Jönsson et al., 2007; Wermelinger et al., 2012; Wermelinger & Seifert, 1998). However, during the Pleistocene, the climate was much harsher even in central and southern Europe (Wright, 1961) and beetles were probably not able to accomplish more than one generation per year. As there are no direct estimates of the mutation rate for bark beetles, we used the *Drosophila melanogaster* mutation rate of 2.8 × 10^−9^ per site per generation (Keightley et al., 2014). If the actual mutation rate is higher than assumed here, the estimated population sizes will be inflated, and vice versa if it is lower. However, even if the estimates are quantitatively different this should not affect the relationship between the estimates obtained for the datasets, nor the comparisons between genetic groups (Nadachowska-Brzyska et al., 2015).

### SFS-based demographic modelling

#### Data preparation

SFS-based demographic inference was performed using a similar approach as defined above, i.e. using four datasets: 1) whole-genome (including all analysed sites), 2) no-inversions (including only inversion-free regions), 3) inversions-only (including only regions affected by large inversions), and 4) no-inversions-small (including only inversion-free regions of size equal to regions analysed in inversion-only dataset). We used polymorphism data generated exactly as described in the section *Sampling, sequencing and data preparation* above and additionally filtered out all sites located within genes (Powell et al., 2021). Based on the NGSadmix results and results of the previous studies (Mayer et al., 2015; Mykhailenko et al., 2024; Stauffer et al., 1999), we decided to restrict our analysis to the least admixed groups: the southern group (populations from Italy, Austria, Germany and the Czech Republic) and the northern group (populations from Norway, Sweden and Finland; see Results and Figure 1). Due to high admixture, Polish and Carpathian populations were excluded from further analysis. We used easySFS (Gutenkunst et al., 2009) to convert the variant call format to SFS and to project population size for each group. Projection to a smaller sample size, together with averaging over all possible resamplings, allows the construction of a complete data matrix with no missing genotypes. As there is a trade-off between the number of individuals retained and the number of sites, we chose a projection value (number of gene copies) for each dataset that maximised the number of available segregating sites within each group, as suggested by Gutenkunst et al. (2009). The projection values were very similar for all four datasets, for the whole-genome the dataset equalled 146 and 124 gene copies for the northern and southern groups, respectively. For each dataset analysed, we obtained an observed two-dimensional site frequency spectrum (2D SFS), which was used to select the best demographic model and estimate its parameters. To allow dating of demographic events, the observed 2D SFS also included monomorphic sites.

#### Tested models

Demographic models were designed based on the results of genetic clustering and differentiation among populations (two main genetic clusters with little admixture between clusters and little within clusters differentiation; see Results for details). We constructed six demographic models (Figure 2). All models assumed ancestral population split into two descendant populations (as suggested by population genetic structure) and estimated ancestral (N_anc_) and descendant population sizes (N_north_ & N_south_) as well as the time of population divergence (T_d_). *Isolation model* (ISO) assumed no gene flow between descendant populations. *Isolation with migration model* (IM) allowed for constant gene flow after population divergence. *Secondary contact model* (SC) assumed time of isolation after the divergence and secondary contact between descendant populations. In both IM and SC models migration rate (m) was symmetrical. Models ISODE, IMDE and SCDE applied the same gene flow regimes as ISO, IM and SC models respectively, but they allowed also for a single post divergence demographic event (DE, i.e. population expansion or contraction). Exact information on parameter number and starting parameter values used are provided in Table 1 and 2 as well as in Figure 2. We used uninformative starting distributions for all parameters.

**Figure 2.**
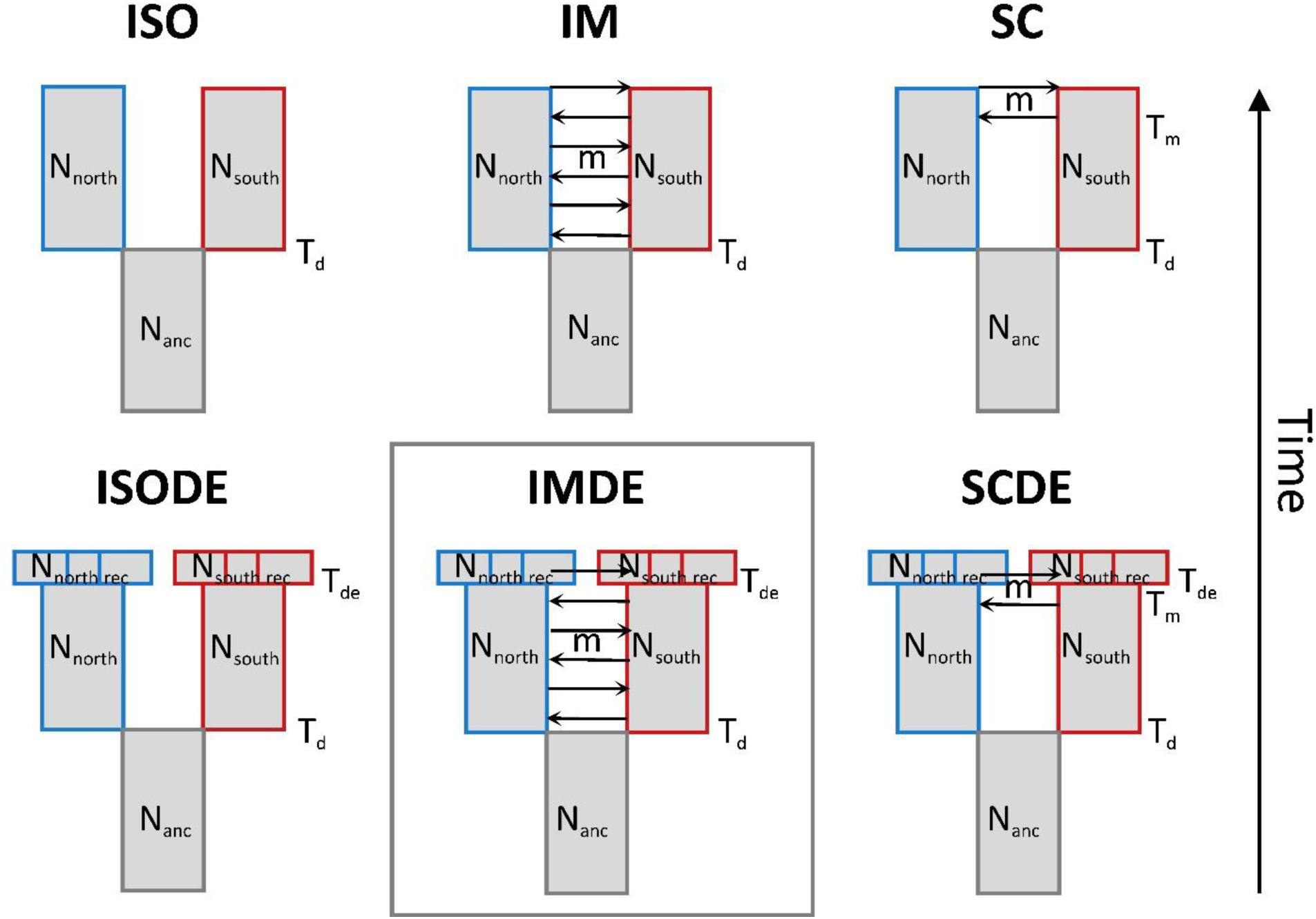
Demographic models investigated. Models were constructed for northern and southern genetic groups. Six divergence scenarios were analysed (ISO - no gene flow, IM - constant gene flow, SC - secondary contact (gene flow after a period of isolation), ISODE - no gene flow plus single instantaneous population size change at any time after divergence, IMDE - constant gene flow plus instantaneous population size change at any time after divergence, SCDE - secondary contact gene flow plus instantaneous population size change at any time after divergence). The frame indicates the best model for all data sets. N_north_, N_south_, N_north rec_, N_south rec_ and N_anc_ - northern, southern, northern after size change, southern after size change and ancestral population sizes, respectively. T_d_ - time of divergence. T_de_ - time of size change (expansion or contraction) in both groups. T_m_ - time of start of migration in secondary contact models. m - migration rate.

**Table 1.** Model fit comparison across datasets. ΔLhood - the difference between the maximum possible value for the likelihood and the maximum likelihood estimated for the best run of a given model, ΔAIC - the difference of Akaike’s information criterion (AIC) between a given model and the best model, Rank - the order of the models determined by ΔAIC, with the best-fitting model ranked 1^st^ and the worst-fitting model ranked 6^th^. Columns contain results for all analysed datasets.

| Model | # of parameters | whole-genome |  |  | no-inversions |  |  | inversions-only |  |  | no-inversions-small |  |  |
| --- | --- | --- | --- | --- | --- | --- | --- | --- | --- | --- | --- | --- | --- |
| | | $\Delta L_{hood}$ | $\Delta AIC$ | Rank | $\Delta L_{hood}$ | $\Delta AIC$ | Rank | $\Delta L_{hood}$ | $\Delta AIC$ | Rank | $\Delta L_{hood}$ | $\Delta AIC$ | Rank |
| ISO | 4 | -28631 | 107371 | 6 | -20629 | 81935 | 6 | -8716 | 26012 | 6 | -9031 | 35020 | 6 |
| IM | 5 | -11927 | 30447 | 4 | -7761 | 22679 | 4 | -4726 | 7636 | 4 | -3466 | 9393 | 4 |
| **SC** | 6 | -9979 | 21479 | 3 | -6721 | 17892 | 3 | -3995 | 4275 | 3 | -3043 | 7446 | 3 |
| **ISODE** | 7 | -24570 | 88675 | 5 | -18164 | 70587 | 5 | -7427 | 20081 | 5 | -7862 | 29641 | 5 |
| **IMDE** | 8 | -5314 | 0 | 1 | -2835 | 0 | 1 | -3066 | 0 | 1 | -1425 | 0 | 1 |
| **SCDE** | 9 | -5468 | 709 | 2 | -3017 | 840 | 2 | -3209 | 660 | 2 | -1509 | 388 | 2 |

#### Model choice and parameter estimation

Simulations, model choice, and parameter estimation were performed using fsc2.7 (Excoffier et al., 2021). We ran maximum likelihood estimation 200 times for all models (to find the global optimum), each time with 50 rounds of parameter optimisation and 200,000 coalescent simulations to approximate the expected 2D SFS. For each model we selected the run that minimised the difference (ΔLhood) between the maximum possible value for the likelihood (MaxObsLhood) and the maximum likelihood estimated (MaxEstLhood). The Akaike information criterion (AIC) was calculated to account for differences in the number of parameters between models. The order of the models was determined by the difference of AIC, i.e. ΔAIC (Excoffier et al., 2021; Marchi et al., 2021; Rougemont et al., 2020). To assess whether the models differed significantly, we ran each model with its best parameter values 100 times and compared the resulting likelihood distributions. If the likelihood ranges of two models overlapped, they were not significantly different, i.e. provide an equally good fit to the observed data. The fit of the models was also visually inspected by comparing the fit of the expected 2D SFS under the models with the observed 2D SFS (Figure 5). The same approach was applied for the marginal 1D SFS (Figure S5-S10). Confidence intervals for the parameters were computed for the best demographic model using the estimates obtained from 100 block-bootstrapped (pseudo-observed) datasets. The parameter estimation process for each pseudo-observed dataset was conducted in the same manner as it was done for the observed dataset, i.e. maximum likelihood estimation was run 200 times, with 50 rounds of parameter optimization and 200,000 coalescent simulations each time.

## Results

### Genome-wide variation and population genetic structure

The whole-genome dataset comprised of over 1.9 million variants after filtering (for the exact dataset sizes see Table S2). All the population genetic summary statistics differed between the inversions-only and no-inversions datasets (p≤0.001). Nucleotide diversity (π) was higher in the regions of the genome affected by inversions (inversions-only) than in the colinear genome (no-inversions, 0.0072 vs 0.0067, respectively), while Tajima’s D was more negative in the collinear genome (−0.86 vs −1.08). Linkage disequilibrium (LD), measured as average LD between SNPs separated by 10,000 base pairs was higher in the regions of the genome affected by inversions (0.079 vs 0.016). Differentiation between the southern and northern populations measured by both *F_ST_* and *d_XY_* was higher in regions of the genome affected by inversions (Table S3). Consistent with previous studies (Mayer et al., 2015; Mykhailenko et al., 2024; Stauffer et al., 1999), the most likely number of genetic clusters was K=2, with populations from the southern and northern parts of the sampling range grouping separately (Figure 1) and individuals from Poland being highly admixed. Apennine and Alpine populations clustered together with German, Austrian and Czech populations (southern group), whereas Carpathian populations showed a moderate degree of admixture between the two main genetic groups (approximately 85% southern ancestry; Figure 1). All pairwise *F_ST_* values were low (0-0.034) between all population comparisons (Table S4).

### Long term effective population size modelling with PSMC

The shapes of the PSMC curves were similar for all analysed individuals (Figure 3, Figure S1) and differed between groups only in the most recent period (<20 kya; Holocene). A recent increase in Ne was observed in the northern and Polish populations, but not in the southern and Carpathian populations (Figure 3, Figure S1). Estimates based on the whole-genome data showed that ancient (0.4-3 Mya, i.e. before the population split) Ne was around 0.2-0.3 million and was relatively stable until 400 kya, when Ne started to increase and reached a maximum of 1-3.5 million around 70-100 kya. Ne then declined to the level of several hundred thousand by 20-30 kya. The Ne estimates based on the collinear parts of the genome (no-inversions) were congruent with the estimates obtained for the whole-genome dataset. However, the estimates obtained from the data containing only inversions (inversions-only) varied widely. First, for some of the individuals, recent (10-20 kya) Ne estimates were increased compared to the estimates for the colinear part of the genome, however the same pattern was present for no-inversions-small dataset. In the more distant time frame (70-100 kya), 55 out of 129 individuals (43%) had effective population size estimates that were substantially (twice or more) higher than those obtained for the colinear genome of the same size (no-inversions-small). To test the association of inversion genotypes with this pattern, estimates for all individuals (regardless of genetic ancestry) obtained from the inversions-only dataset were assigned to a category of higher (>2x), similar (0.5-2x) or lower (<0.5x) than the highest estimate obtained for the no-inversions-small dataset in this time period. In the group with high Ne (>2x), we found deviations from HW equilibrium, most likely driven by the excess of heterozygotes, in the inversions Inv16.1+Inv23.1 (note that Inv16.1 and Inv23.1 are part of the same inversion detected on two separate contigs) and the deficit of heterozygotes in Inv15 (Table S5). In the group that gathers individuals with Ne similar to the no-inversion-small dataset, we found deviations from HW equilibrium in inversions Inv16.1+Inv23.1 and in Inv16.2+Inv23.2. Deviations in Inv16.1+Inv23.1 were most likely driven by the deficit of heterozygotes. However, these result has to be interpreted with caution since after FDR correction none of the deviations remained significant (Table S5). In the group classified as low Ne, we found no deviations from HW equilibrium in any of the analysed inversions, probably due to too low sample size (n=12). In most (78%) of individuals ancient (>1 Mya) Ne estimates based on the inversions-only dataset were slightly inflated compared to other datasets and extended further into the past (up to 4 Mya).

**Figure 3.**
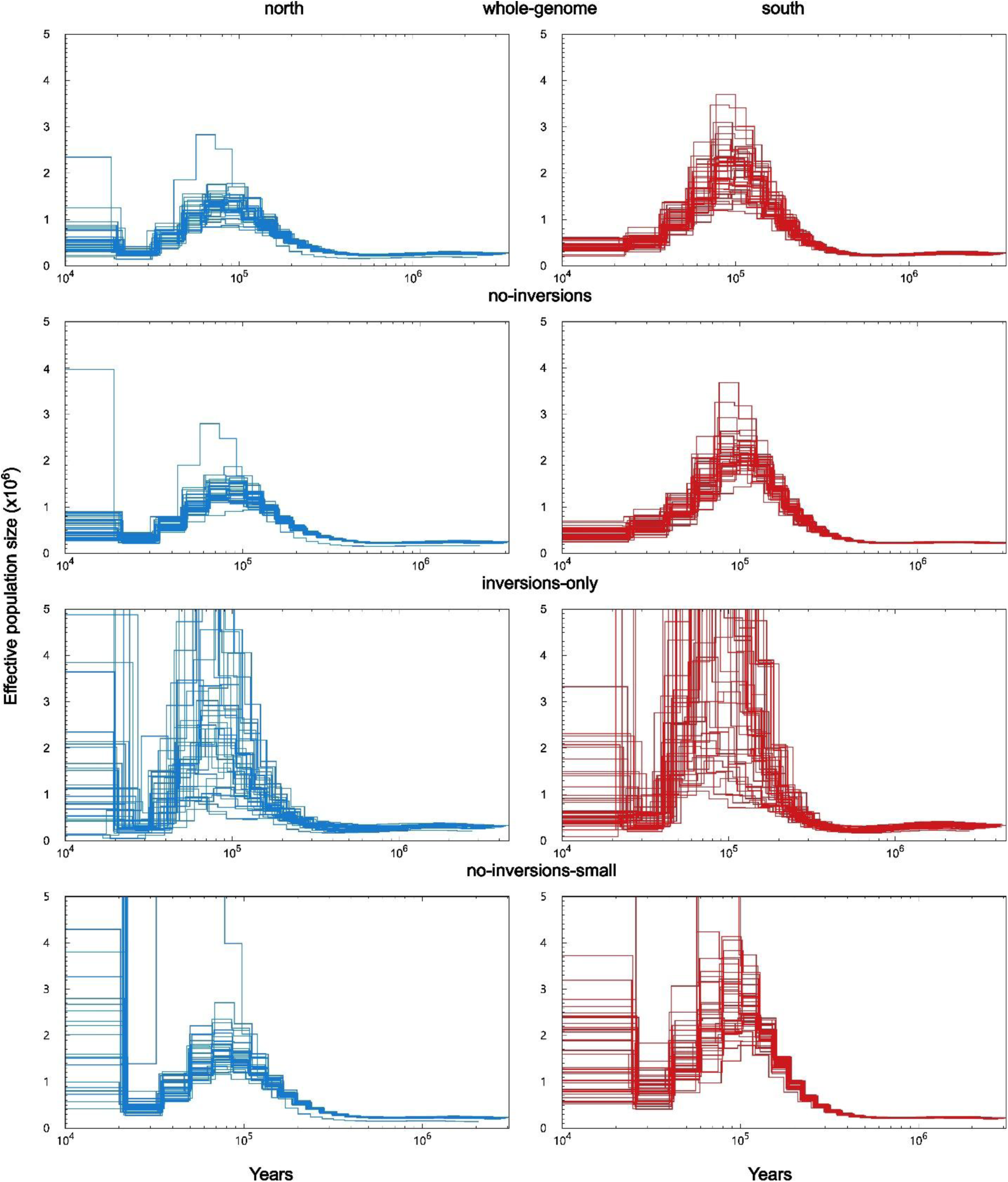
*Pairwise Sequential Markovian Coalescent (PSMC)* analysis results for the northern and southern genetic groups. Rows show results for the four datasets, columns for the two genetic groups. PSMC plots for all individuals assigned to a given group. X-axis - time in years (log scale). Y-axis - population size times 10^6^.

### SFS-based demographic history modelling

We tested six demographic models for each dataset. For each model, we selected the run that minimised the ΔLhood (difference between the maximum possible value for the likelihood and the maximum likelihood estimated). The ranking of the models (based on AIC) was the same regardless of the dataset analysed (Table 1). The best model was IMDE (the model assuming constant post-divergence gene flow and a single size change), while the second-best model was SCDE (secondary contact with a single size change), with a rather small ΔAIC between these two best models (Table 1) and similar expected 2D and 1D SFS under both models (Figure 5, Figure S5-S10). The largely overlapping ΔLhood distributions between IMDE and SCDE suggest that the fit of these two models is comparable (Figure S2), but the non-overlapping likelihood distributions for the best parameter estimates prove that the models differ in their fit to the observed data (Figure S3). The two worst models were those that did not allow for post-divergence gene flow (ISO & ISODE). Overall, the presence of size change allowed the models to perform better than their demographically stable counterparts (Table 1, Figure 5, Figure S2, Figure S5-S10).

Parameter values inferred from the best model (IMDE) for the no-inversions dataset (representing part of the genome least affected by selection) imply a relatively recent (79 kya) group divergence and a very recent (7 kya) demographic expansion in both groups (Table 2, Figure 4). The most recent population size of the southern group was more than 2 times larger than that of the northern group (1.83 million vs. 0.79 million). Before the demographic expansion (7-79 kya), the southern group was more than 4 times larger than the northern group (0.82 million vs. 0.18 million). Thus, the southern group expanded after the population split, as the ancestral population size (for both groups) was estimated to be 0.25 million. The estimated migration rate of 1.6 × 10^−5^, translated into an effective number of 19 migrants per generation (Nm), suggests a constant and relatively high level of gene flow between the groups, with migration from south to north being almost 4 times higher than vice versa (Figure 4, Table 2).

**Figure 4.**
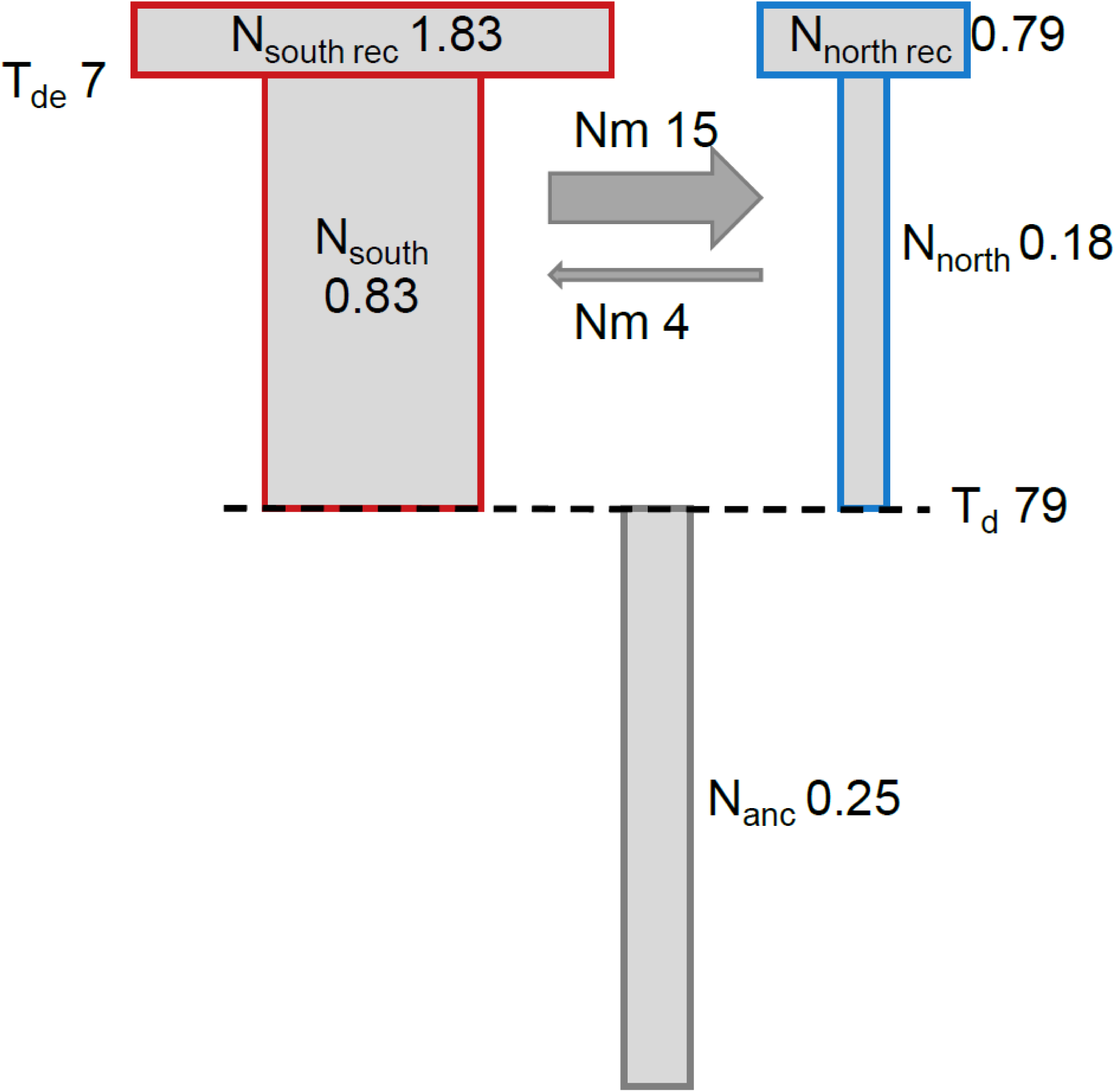
Best demographic model (IMDE) and parameter estimates obtained for the no-inversions dataset. Population sizes given in millions. Time estimates given in thousands years ago.

**Table 2.** Parameter estimates for the best demographic model i.e. IMDE, for all datasets. ML - maximum likelihood. Minimum and Maximum – starting ranges for maximum likelihood estimation. N_north_, N_south_, N_north rec_, N_south rec_ and N_anc_ - northern, southern, northern after size change, southern after size change and ancestral population sizes, respectively. T_d_ - time of divergence. T_de_ - time of size change (expansion or contraction) in both groups. m - migration rate.

| Parameter | Minimum | Maximum | ML estimates (95% CI) |  |  |  |
| --- | --- | --- | --- | --- | --- | --- |
|  |  |  | whole-genome | no-inversions | inversions-only | no-inversions-small |
| $N_{\text{north rec}}^2$ | 0.005 | 0.5 | 0.80 (0.67-1.08) | 0.79 (0.68-1.11) | 0.67 (0.60-0.93) | 0.68 (0.59-1.11) |
| $N_{\text{south rec}}^2$ | 0.005 | 0.5 | 1.70 (1.31-1.98) | 1.83 (1.25-1.93) | 1.47 (1.21-1.93) | 2.17 (1.17-1.93) |
| $N_{\text{north}}^2$ | $0.1 * N_{\text{north rec}}$ | $10 * N_{\text{north rec}}$ | 0.19 (0.18-0.20) | 0.18 (0.17-0.18) | 0.20 (0.20-0.27) | 0.18 (0.17-0.19) |
| $N_{\text{south}}^2$ | $0.1 * N_{\text{south rec}}$ | $10 * N_{\text{south rec}}$ | 0.82 (0.76-0.99) | 0.83 (0.81-1.00) | 0.82 (0.68-0.98) | 0.91 (0.87-1.13) |
| $N_{\text{anc}}^2$ | 0.005 | 0.5 | 0.26 (0.26-0.27) | 0.25 (0.24-0.25) | 0.30 (0.28-0.30) | 0.28 (0.27-0.30) |
| $T_d^1$ | 1 | 1000 | 73 (70-80) | 79 (76-86) | 61 (58-70) | 84 (78-98) |
| $T_{\text{de}}^1$ | 0.001 | $T_d$ | 7.65 (6.15-8.58) | 7.00 (5.61-7.92) | 9.21 (6.50-10.87) | 8.03 (5.72-9.44) |
| $m^3$ | $1 * 10^{-8}$ | $1 * 10^{-4}$ | 1.42 (1.39-1.55) | 1.60 (1.52-1.72) | 1.26 (1.12-1.37) | 1.56 (1.44-1.72) |

<sup>1</sup> time given in thousands of generations.
<sup>2</sup> population size given in millions.
<sup>3</sup> migration rate estimates given in $10^{-5}$

The demographic parameters inferred for the best model (IMDE) were generally similar between the datasets. However, there were some differences when comparing the estimated parameter values. Notably, the Ne estimates for the southern group obtained from the no-inversions-small dataset were higher than those obtained from all the other datasets (Table 2). This is most likely due to the accidental enrichment of low-frequency variants in the southern group during the random sampling of the genomic regions (Figure 5). Therefore, we focus more on interpreting the results obtained for the other three datasets. First, the divergence time (T_d_) was more recent (23%) for the inversions-only dataset (61 kya and 79 kya, for inversions-only and no-inversions datasets, respectively), whereas the estimate for the whole-genome dataset was between that obtained with the two other datasets (73 kya). Second, the ancestral population size (N_anc_) was 19% larger for the inversions-only dataset than for the no-inversions dataset (0.30 vs. 0.25 million, respectively), with the estimate for the whole-genome dataset being intermediate (0.26 million). Third, the most recent population size estimates (N_north rec_ & N_south rec_) derived from the inversion-only dataset were lower for both groups (15% and 19% for northern and southern populations, respectively) than those derived from the no-inversions dataset. The most recent population size estimates obtained for the whole-genome dataset were intermediate for the southern group, but not for the northern group, where they were almost identical to those from the no-inversions dataset (Table 2). The pre-demographic expansion population size for the southern group (N_south_) was almost identical for the three datasets. For the northern group, the population size estimate between divergence and demographic event (N_north_) was 12% higher for the inversions-only dataset. Fourth, the migration rate estimates (m) were lower (21%) for the inversions-only dataset compared to the no-inversion dataset, with an intermediate value estimated for the whole-genome dataset. Finally, the estimate of the time of the demographic event (T_de_) in the descendant populations was 32% older in the inversions-only dataset than in the no-inversions dataset (9.2 kya vs. 7 kya), again with the intermediate genome-wide estimate (7.7 kya).

**Figure 5.**
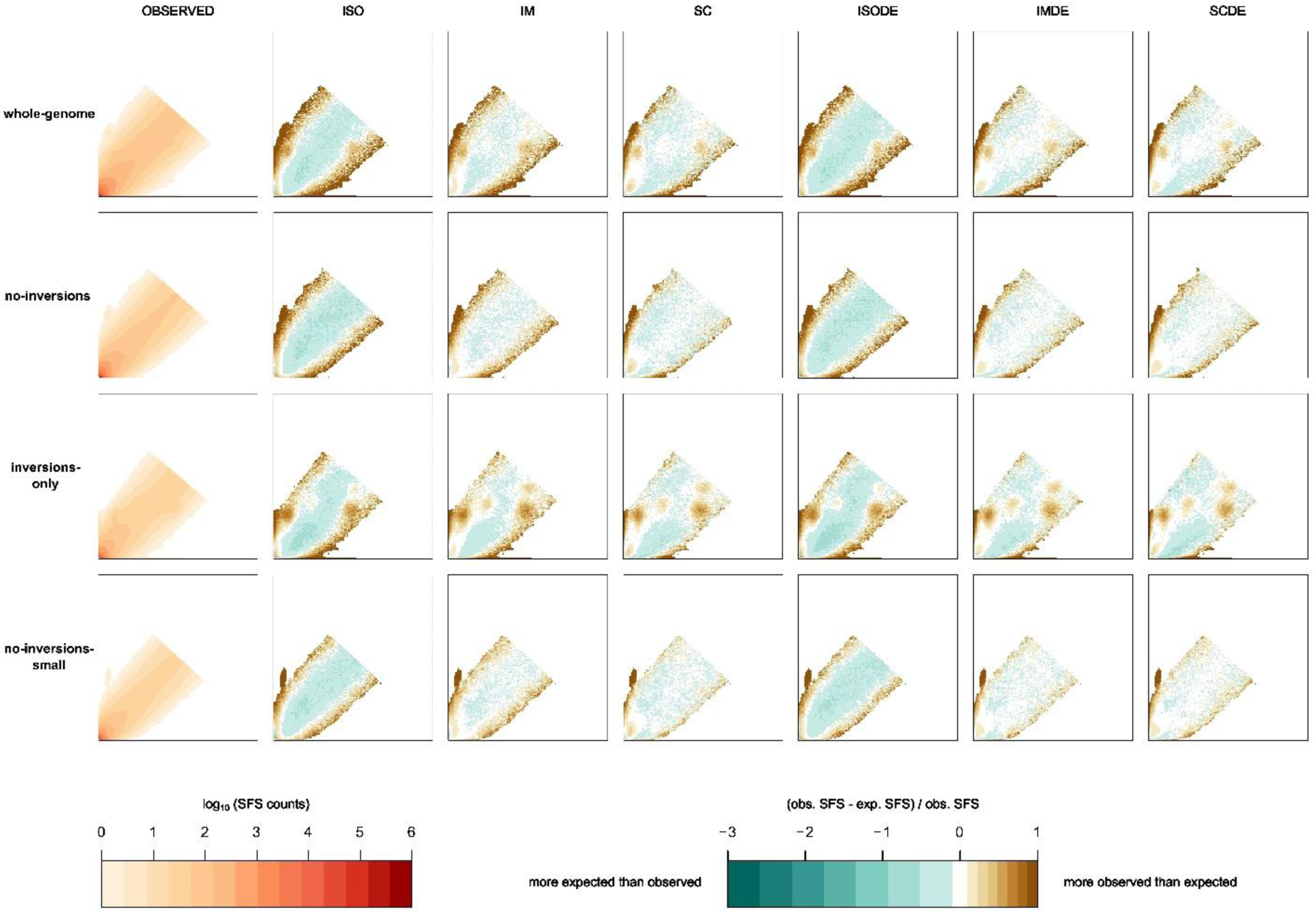
Comparison of observed and simulated 2D SFS. The first column shows the observed 2D SFS for each dataset. Further columns compare the fit of the expected 2D SFS under the given model to the observed data. For each 2D SFS, the northern and southern groups are on the y-axis and x-axis, respectively. We present 2D SFS entries used to calculate the likelihood, i.e. with at least one polymorphism.

## Discussion

### Genome-wide genetic structure in the context of previous findings

Consistent with previous reports based on a few genetic markers (Bertheau et al., 2013; Ellerstrand et al., 2022; Mayer et al., 2015; Stauffer et al., 1999) and the recent whole-genome re-sequencing study (Mykhailenko et al., 2024), *I. typographus* was characterised by very weak genetic differentiation and structuring into two genetic clusters (northern and southern). This result is consistent with recent divergence and high levels of gene flow, which is not surprising for a species with a high dispersal capacity (Inward et al., 2024; Nilssen, 1984). We found that all Carpathian populations were almost equally admixed (84-85% southern ancestry), suggesting an admixture event within the Carpathians or recent colonisation of the Carpathians by an already admixed population. This picture differed from that observed in Poland, where population admixture levels varied widely over relatively short distances (17-63% southern ancestry), suggesting a transition zone between the two genetic clusters in Poland. Which is consistent with the results of Mykhailenko et al. (2024) who showed the highest nucleotide diversity in the Polish populations. The overall genetic structure of the spruce bark beetle is similar to the genetic structure of its host plant (Chen et al., 2019; Tollefsrud et al., 2008). However, Norway spruce is characterised by the presence of three major genetic groups: northern (Fennoscandian) and two smaller southern domains (Alps and Carpathians).

### Historical demography

Our results revealed a Late Pleistocene (79 kya, within the Last Glacial Period) divergence between the southern and northern groups (Figure 4, Table 2), suggesting isolation in two distinct glacial refugia. This is consistent with the existence of separate refugia for Norway spruce at that time (Lagercrantz & Ryman, 1990; Ravazzi, 2002; Tollefsrud et al., 2008). Both methods used to reconstruct historical demography suggested population growth around the time of divergence, consistent with either true Ne changes or geographic subdivision at that time (Li & Durbin, 2011; Mather et al., 2020; Mazet et al., 2015; Nadachowska-Brzyska et al., 2016).

The inferred post-divergence demographic expansion appears to be stronger in the southern group than in the northern group. According to the SFS-based model, the southern group experienced a substantial (3-fold) population expansion immediately after divergence, while the northern group remained stable. The expansion of the southern group after the population split could be interpreted as a result of population substructure due to subdivision into different refugia during glacial-interglacial cycles (refugia within refugia), as spruce is known to have survived the Last Glacial Maximum (20 kya – 26 kya), in several southern refugia (Ravazzi, 2002; Tollefsrud et al., 2008). Although there were some slightly elevated *F_ST_* values between some populations from the southern group (Table S4), we found no support for substructure when examining admixture proportions estimated for higher numbers of genetic groups (K = 3 and K = 4; Figure S4). The growth of the southern population was supported by the wide and undivided distribution of Norway spruce in central and southeastern Europe during 110-34 kya (Ravazzi, 2002). Therefore, we tend to interpret increased post divergence Ne estimates as a signal of real growth of the southern group during the Last Glacial Period.

According to the SFS-based model, the Ne of both groups increased during the Holocene (7 kya), but this time the expansion was stronger in the northern group (4-fold) than in the southern (2-fold). This result is supported by the PSMC approach, which also inferred an expansion of the northern group within the last 20 ky. However, as PSMC is not optimal for estimating Ne in periods younger than 20 kya (Li & Durbin, 2011), we focused on the estimates provided by the SFS-based approach for these more recent periods. The demographic expansion in the Holocene is consistent with some previous mtDNA-based inferences (Bertheau et al., 2013), but not with others that suggest a relatively stable population size over time, except for a very recent increase (<100 years) associated with human spruce cultivation (Mayer et al., 2015). It has been shown that Norway spruce expanded from southern refugia at the beginning of the Holocene, reconnecting western European refugia by 9 kya (Tollefsrud et al., 2008). Expansion across the northern Norway spruce range from a single refugium located in the Russian Plain was completed by 6 kya, with both populations coming into present-day secondary contact in Poland (Tollefsrud et al., 2008), but see Chen et al., (2019), who suggest contraction of all Norway spruce groups at the beginning of the Holocene. Therefore, our results showing demographic expansion of both groups around 7 kya are congruent with the natural expansion of Norway spruce, the main host plant species of *I. typographus*.

Taking all population size changes together, the post-divergence expansion of the southern group was greater than that of the northern group (mainly due to a substantial intra-Pleistocene expansion immediately after the ancestral population split, i.e. 79 kya), and suggest that the southern group is more than twice as large as the northern group (1.8 vs. 0.8 million). This result is consistent with the more negative Taijma’s D value inferred for the southern group (Mykhailenko et al., 2024). Given the larger contemporary range of spruce in the northern group compared to the southern group (Caudullo et al., 2017), such a discrepancy in Ne may be due to bark beetle populations having more generations per year and more frequent outbreaks in the southern part of the species’ range, resulting in a temporal increase in population size by several orders of magnitude. Although we cannot disentangle these two scenarios with the present data, we believe that they may be strongly linked, as the ability to produce 2-3 generations per year may facilitate outbreaks in southern populations. Importantly, as climate change continues, populations in the northern group are also predicted to be able to achieve 2 generations per year facilitating their propensity for outbreaks and increasing effective population size (Gohli et al., 2024; Jaime et al., 2024; Jakoby et al., 2019; Jönsson et al., 2009).

### Demographic history of species and influence of inversions on inference

Observed genome-wide variation is the result of different evolutionary forces including drift and selection acting within populations/species over long evolutionary timescales. However, disentangling their effects on genomic patterns is often challenging (Charlesworth et al., 1993; Jensen et al., 2005; Johri et al., 2021; Li et al., 2012; Soni et al., 2024). One of the difficulties arises from the fact that many possible evolutionary scenarios produce similar patterns within the genome, and the other arises from the difficulties associated with detecting regions under selection. A good example are chromosomal inversions. Inversions were one of the first polymorphisms described in natural populations (Sturtevant, 1926; Sturtevant, 1917) and since then, have been detected in many organisms, including *Drosophila* (Graubard, 1932), mosquitoes (Coluzzi et al., 1977), sunflowers (Burke et al., 2004), snails (Koch et al., 2021), and many more (Wellenreuther & Bernatchez, 2018). Failure to account for inversion polymorphisms has also been shown to bias estimates of population structure (Corbett-Detig & Hartl, 2012; McKinney et al., 2020; Seich al Basatena et al., 2013) and the results of selection scans (Berdan et al., 2023; Lotterhos, 2019), or to miss sex-determining loci (McKinney et al., 2020).

With novel genomic developments and the accumulation of population genomic data, we are witnessing a recent surge in the discovery of complex inversion polymorphism landscapes (Huang et al., 2020; Matschiner et al., 2022; Meyer et al., 2024; Mykhailenko et al., 2024; Mérot et al., 2021; Porubsky et al., 2022; Reeve et al., 2023). Since many polymorphic inversions under different evolutionary pressures (Nosil et al., 2023) are often present within a single genome, many researchers face the challenge of analysing inversion-rich genomes. Therefore, we need a much better understanding of the influence of complex genomic inversion architecture on evolutionary inference.

In our previous study (Mykhailenko et al., 2024), we identified 27 large (0.1-10.8 Mb) inversions, representing a substantial fraction of the genome (approximately 28%). Most of the inversions are polymorphic across the species range and appear to be 2 orders of magnitude older than the estimated divergence time between groups (age of inversions: 0.5-2.6 My vs. north-south divergence: 79 ky). Importantly, many of these are present at high frequencies and are likely to be maintained by balancing selection (Mykhailenko et al., 2024). Such a scenario has great potential to bias demographic inferences (Novo et al., 2023). Moreover, even if recombination reduction is caused by factors other than inversions, it might affect genetic variation similarly (Berdan et al., 2023; Charlesworth, 2012; Corbett-Detig et al., 2015). Therefore, regardless of the reason, a potential effects of linkage should not be overlooked when inferring population history (Johri et al., 2021). To test the influence of inversions on demographic inference, we compared the results obtained for the whole-genome, the colinear part of the genome (no-inversions), and the part of the genome restricted to inversions (inversions-only), using and evaluating two widely used methods: PSMC and SFS-based modelling. Our results indicate that both methods are affected to some extent by the presence of inversions, but in different ways.

Very similar PSMC curves were obtained for all the datasets analysed, regardless of whether we included or excluded inversion regions. Therefore, it appears that even if a substantial fraction of the genome is affected by the presence of inversions, this does not affect the accuracy of PSMC inference. However, when data are restricted to inversion regions, deviations from the true demographic scenario can be substantial and differ between individuals depending on the inversion genotypes they carry. This may be due to a reduced ability to correctly estimate temporal Ne changes due to the smaller data size i.e. reduction to the total length of inversion regions only (here approx. 45Mb)(Gower et al., 2018; Liu & Hansen, 2017; Mather et al., 2020). However, the significant (0.005, but 0.057 after FDR correction) deviation from HW equilibrium found in the group of individuals characterised by the most biased PSMC curves in respect to the estimates obtained from the colinear dataset trimmed to the same size (no-inversions-small), suggests that, in extreme cases, the inferred demographic history may depend on the particular inversion genotype.

In the case of SFS-based modelling, the likelihood of bias due to a single inversion genotype is much lower, as typically many individuals are sampled per population/group, so that the effect (e.g. the magnitude of recombination suppression) of a particular inversion locus depends on its frequency (Seich al Basatena et al., 2013). Nevertheless, the presence of polymorphic inversions evolving under long-term balancing selection may shift the SFS towards an equilibrium/intermediate frequency and an excess of polymorphic sites (Bitarello et al., 2023; Bitarello et al., 2018), which in turn may affect demographic inference. In our study, the same demographic model was chosen regardless of the dataset used to provide the observed 2D SFS. The estimated parameters differed by an average of 18% between the inversions-only and no-inversions datasets. The dataset restricted to inversion regions (inversions-only) gave the highest ancestral Ne estimates and the youngest divergence time between the southern and northern groups. A shift in divergence time towards younger estimates and larger ancestral population sizes was expected, as evolution under balancing selection is known to delay lineage sorting relative to a neutrally evolving genome (Charlesworth & Charlesworth, 2010; Fijarczyk & Babik, 2015; Mailund et al., 2014; Meyer & Thomson, 2001; Wiuf et al., 2004). In the case of recent Ne estimates, we found the opposite effect - the inversions-only dataset provided lower estimates than the dataset without inversions, again consistent with the excess of frequent polymorphisms due to evolution under balancing selection (Bitarello et al., 2018; Charlesworth & Charlesworth, 2010; Fijarczyk & Babik, 2015). Estimates based on the whole-genome dataset and the no-inversions dataset were very similar (about 6% difference between the datasets). On the other hand, considering that the effect of inversions can extend beyond the inverted regions (Adrion et al., 2020; Koury, 2023; Kulathinal et al., 2009; Machado et al., 2007; Navarro et al., 1997; Stevison et al., 2011; Sánchez-Gracia & Rozas, 2011), we can assume that this might be the lower limit of the real difference between the estimates. Thus, polymorphic inversions affect the observed SFS (see Figure 5) and the demographic inference based on it, but these effects are nuanced and in most cases, at least in our study system, affect the precision of the parameter estimates (Table 2). Similar conclusions were recently reached for *Littorina* snails, showing that although the site frequency spectra of datasets including and excluding inversions differed, this did not change the model choice and provided similar split time estimates (Le Moan et al., 2024). It should be noted that the inversions-only dataset represents a mixture of the evolutionary histories of more than two dozen independent or nearly independent genomic regions, which may have been affected to varying degrees by different evolutionary forces (Kapun & Flatt, 2019). Therefore, it would be valuable to evaluate their separate evolutionary histories (Poikela et al., 2024). Approaches based on Ancestral recombination graph (ARG) reconstruction could be particularly helpful in this respect, as they offer computational efficiency and higher accuracy than traditional methods (Lewanski et al., 2024; Nielsen et al., 2025).

It is interesting that both methods appear to be quite robust to potential biases caused by the presence of many intermediate frequency (close to 0.5), relatively old and large inversions. We hypothesise that this may be explained by still effective recombination within the homozygotes (Keightley & Otto, 2006; Martin et al., 2006; Otto & Barton, 2001) and is consistent with the lack of evidence for increased mutation load within the inverted regions (Mykhailenko et al., 2024).

## Conclusions and future directions

Genome-wide analysis revealed two genetic clusters in the spruce bark beetle and a weak genetic differentiation among the populations studied. Demographic history analysis suggested a Pleistocene divergence between the southern and northern groups, probably due to isolation in separate glacial refugia. Both PSMC- and SFS-based modelling suggested a stronger post-divergence population expansion of the southern group. Both methods also supported expansion of the northern group during the Holocene, but the recent effective population size of the southern group was twice that of the northern group. Differences in parameter estimates in different data sets (restricted to inversions and whole-genome data sets with and without inversions) followed theoretical expectations, but the differences found were relatively small.

Despite these results, it is important to emphasise that the impact of inversions may differ between species, depending on the properties of the inversions (size, frequency, evolutionary forces acting on them), the effective population size of the species and its demographic history. The growing accumulation of inversion-rich data provides an exciting opportunity to study the evolutionary history of inversions (Poikela et al., 2024), but also their implications for evolutionary inference. While we should maintain a gold standard to exclude sequences that do not evolve neutrally in demographic inference, we should not miss the opportunity to treat inversions as powerful modifiers of evolutionary history (Berdan et al., 2023) and investigate their influence on diverse aspects of evolutionary biology.

## Acknowledgments

We thank the members of the Genomics and Experimental Evolution Group at the Jagiellonian University for their help in improving this manuscript. This work was funded by the Polish National Science Center 2018/30/E/NZ8/00105 grant to K.N.B. We gratefully acknowledge Polish high-performance computing infrastructure PLGrid (HPC Center: ACK Cyfronet AGH) for providing computer facilities and support within computational grant no. PLG/2025/018231.

## Data Availability

All DNA sequences have been deposited in the National Centre for Biotechnology Information Sequence Read Archive under BioProject ID PRJNA1013983. Scripts used for data processing, generation of intermediate files, and demographic analyses, along with key intermediate datasets, have been archived and are publicly available in the Dryad Digital Repository entry [will be added as soon as possible].

## Author contributions

P.Z. and K.N.B. designed the study and wrote the manuscript. P.Z. performed the main analyses and supervised the processing of the raw data. J.M.G. isolated DNA, prepared samples for sequencing, processed raw sequencing data and performed genetic structure analysis. M.S and M.L.D provided samples. All authors read and approved the final version of the manuscript.

