## Supplementary information for "Demographic history inferred from an inversion-rich spruce bark beetle genome"

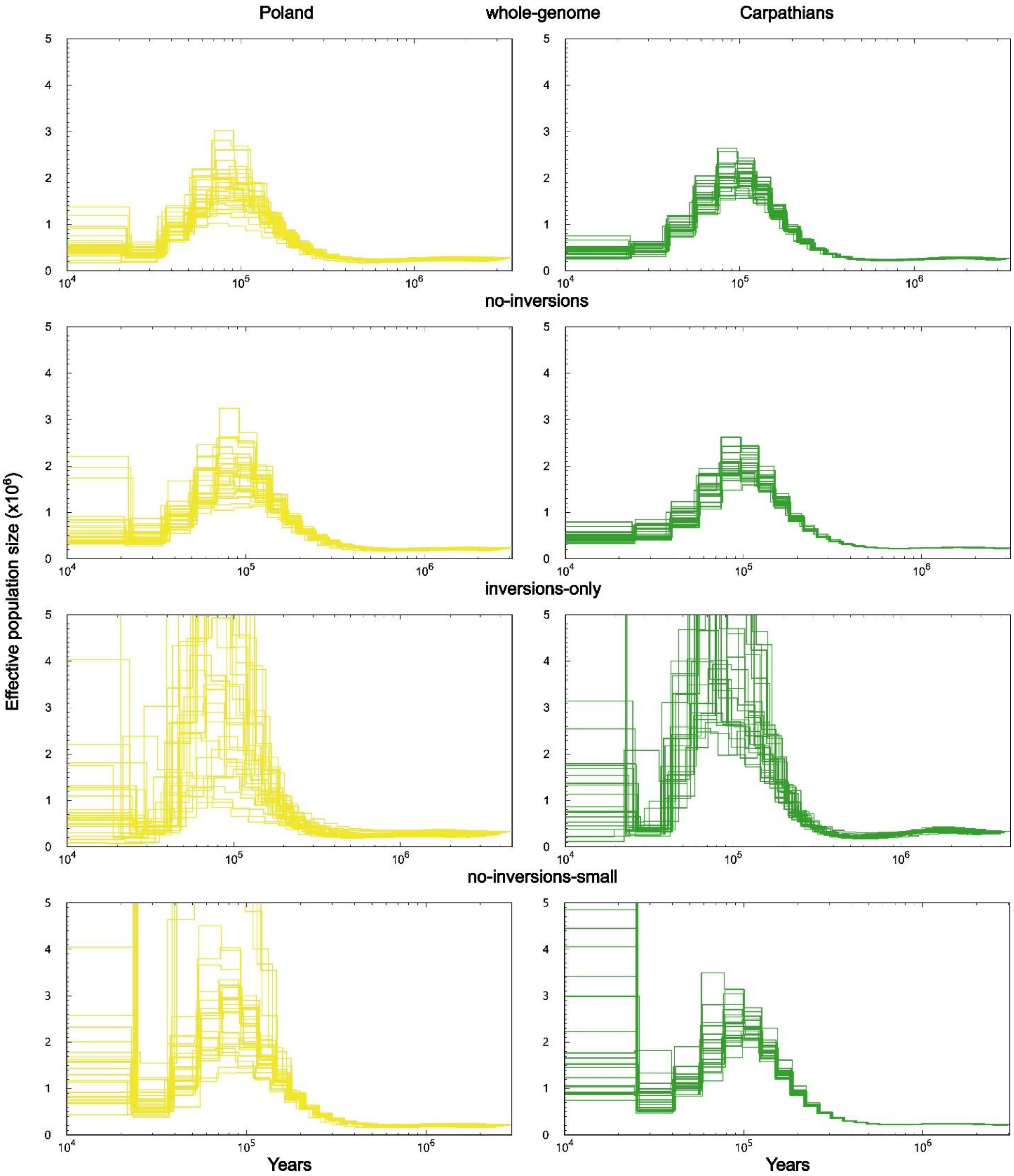


Figure S1. *Pairwise Sequential Markovian Coalescent (PSMC)* analysis results. Rows show results for the four datasets, columns for the two genetic groups (Polish and Carpathian populations). PSMC plots for all individuals assigned to a given group. X-axis - time in years (log scale). Y-axis – population size times $10^6$.

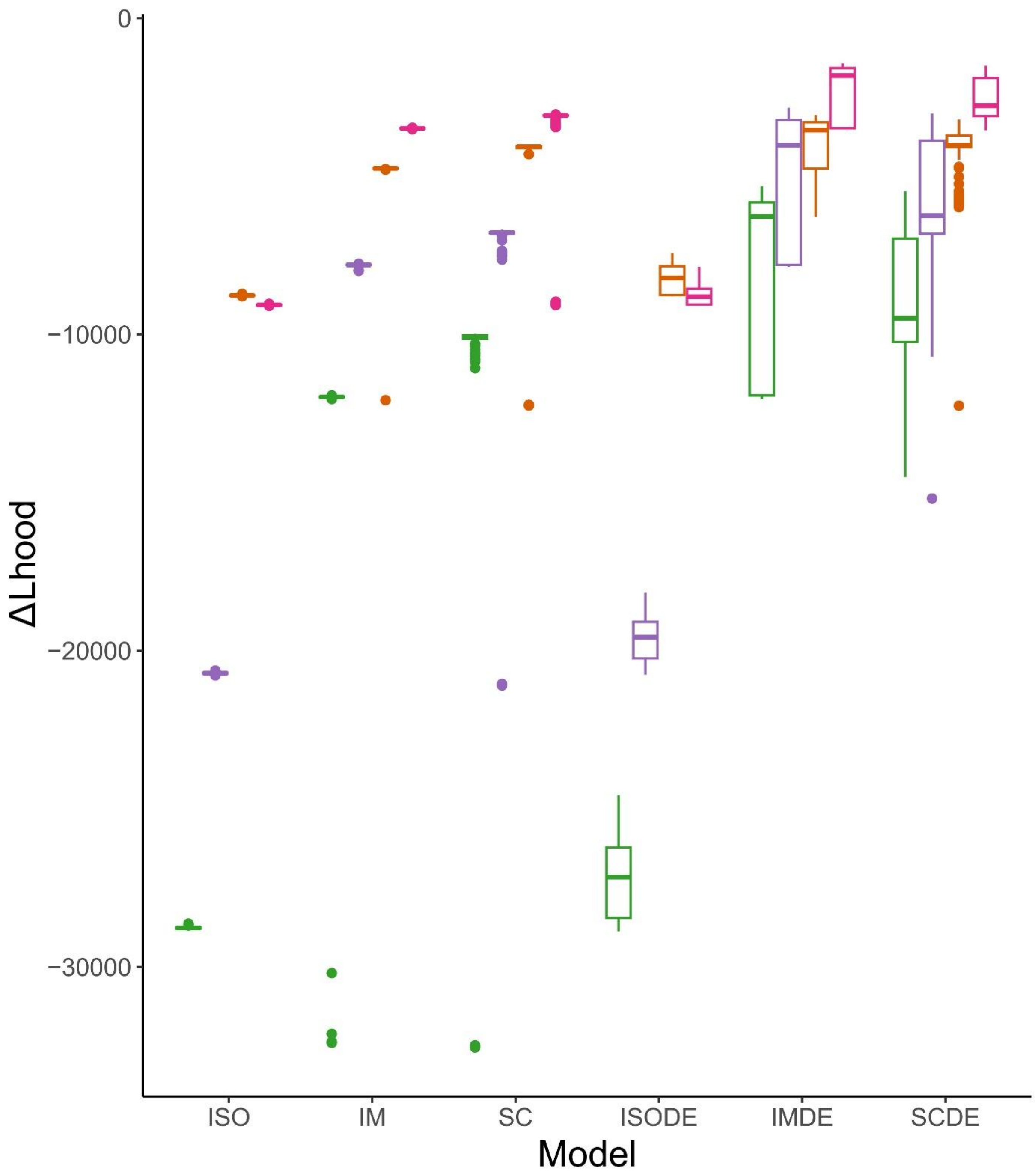


Figure S2. The difference between the maximum possible value for the likelihood and the estimated maximum likelihood (ΔLhood) for each model for 200 runs. Colours show results for the datasets: green - whole-genome, purple – no-inversions, orange – inversions-only, pink – no-inversions-small. The lower and upper hinges correspond to the first and third quartiles. The upper whisker extends from the hinge to the largest value no further than 1.5 times the interquartile range (distance between the first and third quartiles). The lower whisker extends from the hinge to the smallest value no further than 1.5 times the interquartile range of the hinge. Data beyond the end of the whiskers are outliers and are plotted individually.

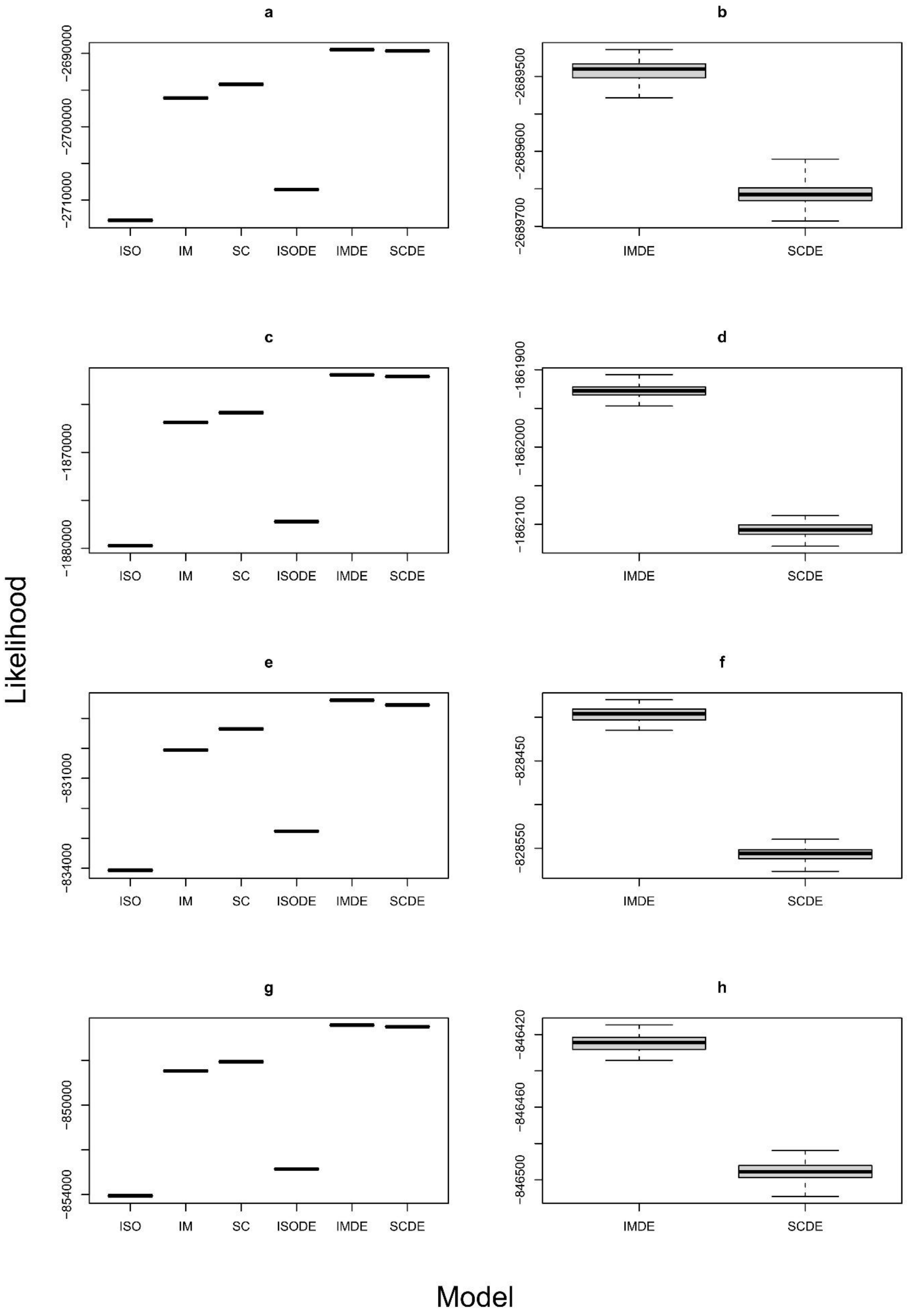
a
−2690000
−2700000
−2710000
ISO
IM
SC
ISODE
IMDE
SCDE
b
−2689500
−2689600
−2689700
IMDE
SCDE
c
−1870000
−1880000
ISO
IM
SC
ISODE
IMDE
SCDE
d
−1861900
−1862000
−1862100
IMDE
SCDE
e
−831000
−834000
ISO
IM
SC
ISODE
IMDE
SCDE
f
−828450
−828550
IMDE
SCDE
g
−850000
−854000
ISO
IM
SC
ISODE
IMDE
SCDE
h
−846420
−846460
−846500
IMDE
SCDE
Likelihood
Model

Figure S3. Model comparison using likelihood distributions. The likelihood provides information about the fit of the model to the observed data for the best parameter estimates. Panels a, c, e, g show likelihoods for all models, while panels b, d, f, h show likelihoods only for the first and second best models, i.e. IMDE and SCDE, respectively (abbreviations explained in *Materials and methods*). Panels a and b show likelihoods for the whole-genome data set, c and d for the no-inversions data set, e and f for the inversions-only data set, g and h for the no-inversions-small data set. Hinges are the first and third quartile. The upper whisker extends from the hinge to the largest value no further than 1.5 times the interquartile range (distance between the first and third quartiles). The lower whisker extends from the hinge to the smallest value at most 1.5 times the interquartile range of the hinge.

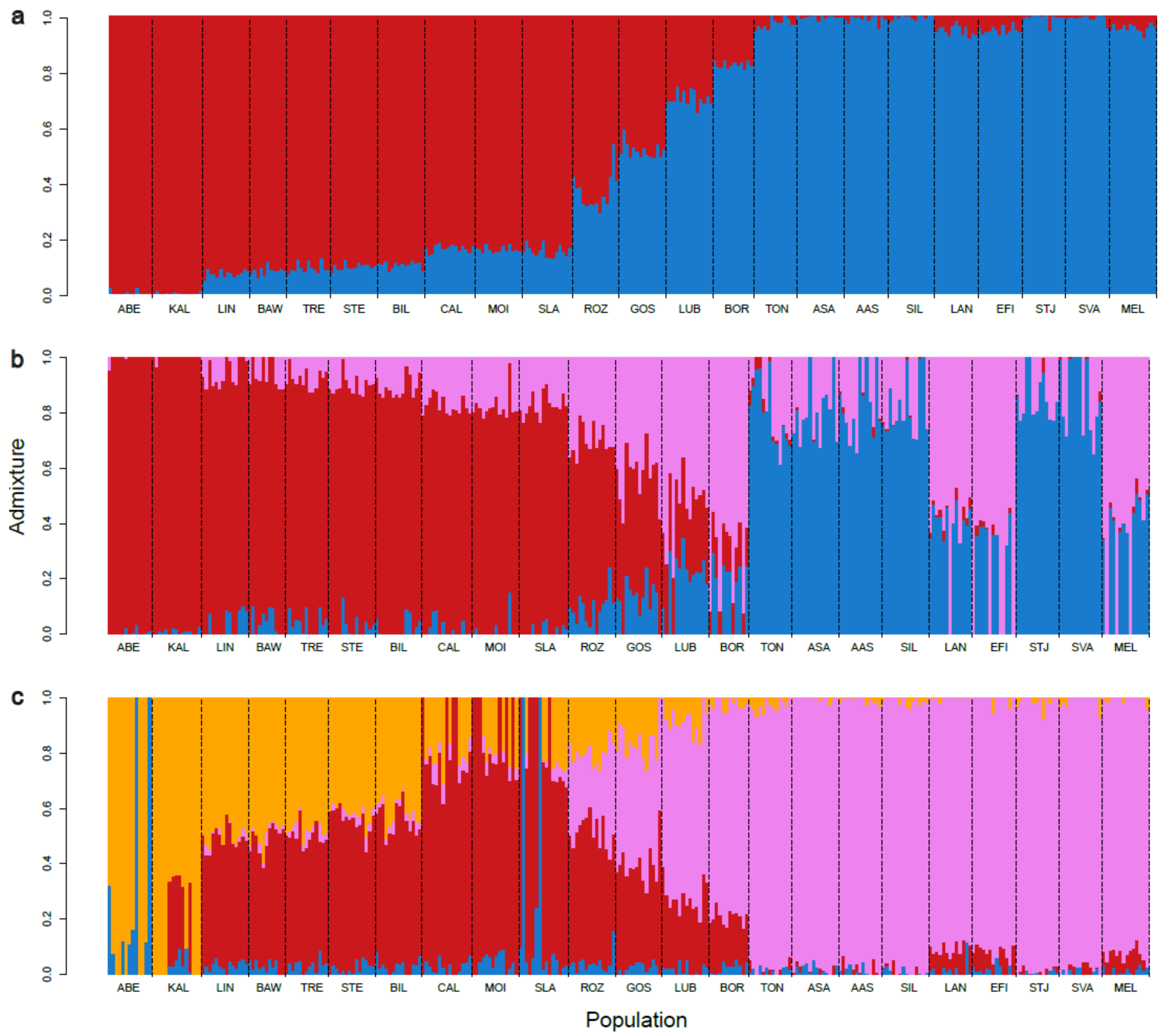


Figure S4. Admixture proportions among *Ips typographus* populations. Individual admixture proportions for different numbers of genetic groups, i.e. a) K=2, b) K=3 and c) K=4. Population abbreviations as assigned in Table S1.

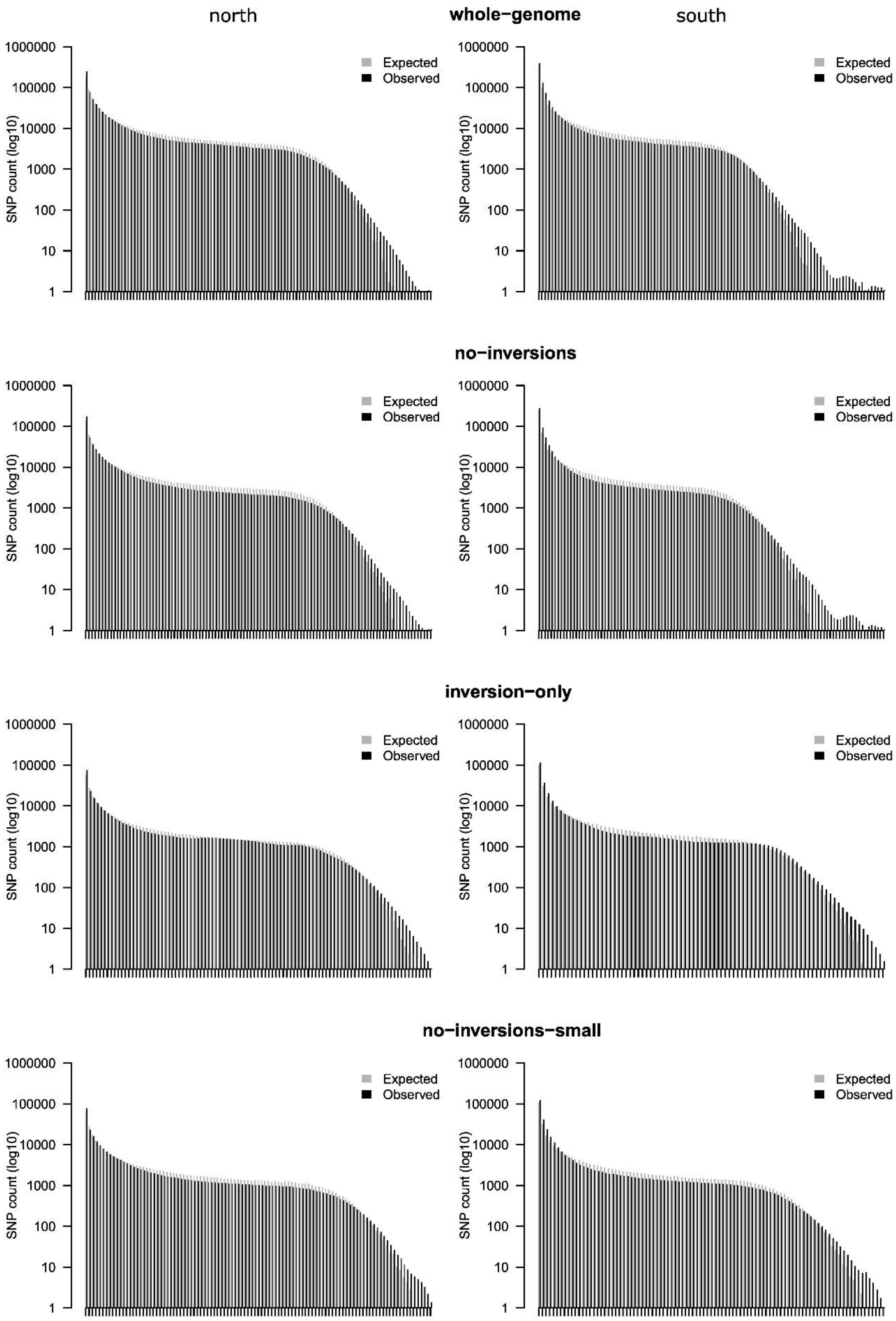
north
whole−genome
south
no−inversions
inversion−only
no−inversions−small
SNP count (log10)
1000000
100000
10000
1000
100
10
1
Expected
Observed

Figure S5. Expected one dimensional site frequency spectra (1D SFS), generated under the ISO demographic model, compared with observed 1D SFS across four genomic datasets: whole-genome, no-inversions, inversions-only, and no-inversions-small. Each dataset is represented by two panels for northern (left) and southern (right) populations. Bars show SNP counts per allele frequency bin; only bins with at least one count are shown. Observed values are shown in black and expected values in grey.

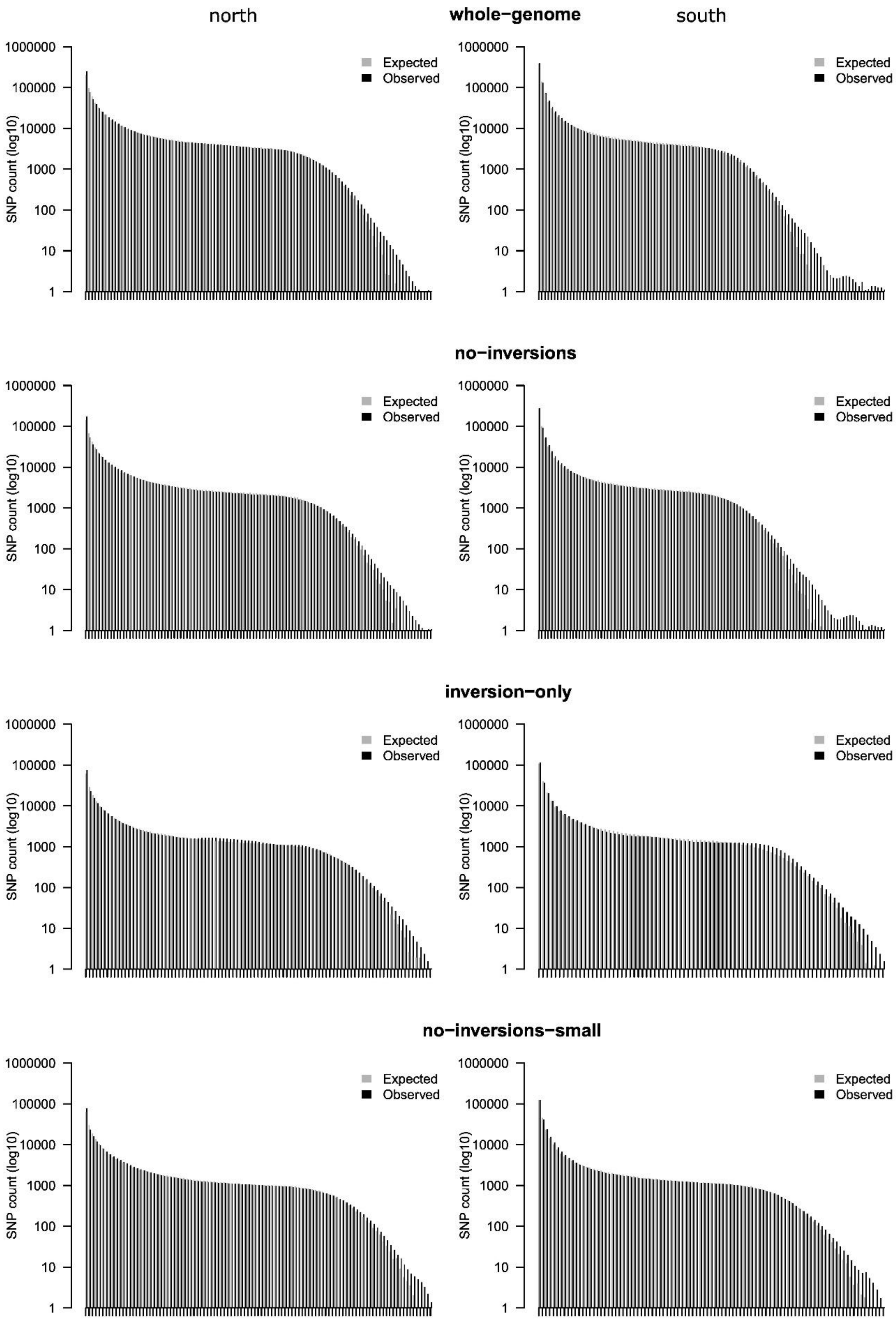

north
whole−genome
south
no−inversions
inversion−only
no−inversions−small
Expected
Observed
SNP count (log10)
1000000
100000
10000
1000
100
10
1

Figure S6. Expected one dimensional site frequency spectra (1D SFS), generated under the IM demographic model, compared with observed 1D SFS across four genomic datasets: whole-genome, no-inversions, inversions-only, and no-inversions-small. Each dataset is represented by two panels for northern (left) and southern (right) populations. Bars show SNP counts per allele frequency bin; only bins with at least one count are shown. Observed values are shown in black and expected values in grey.

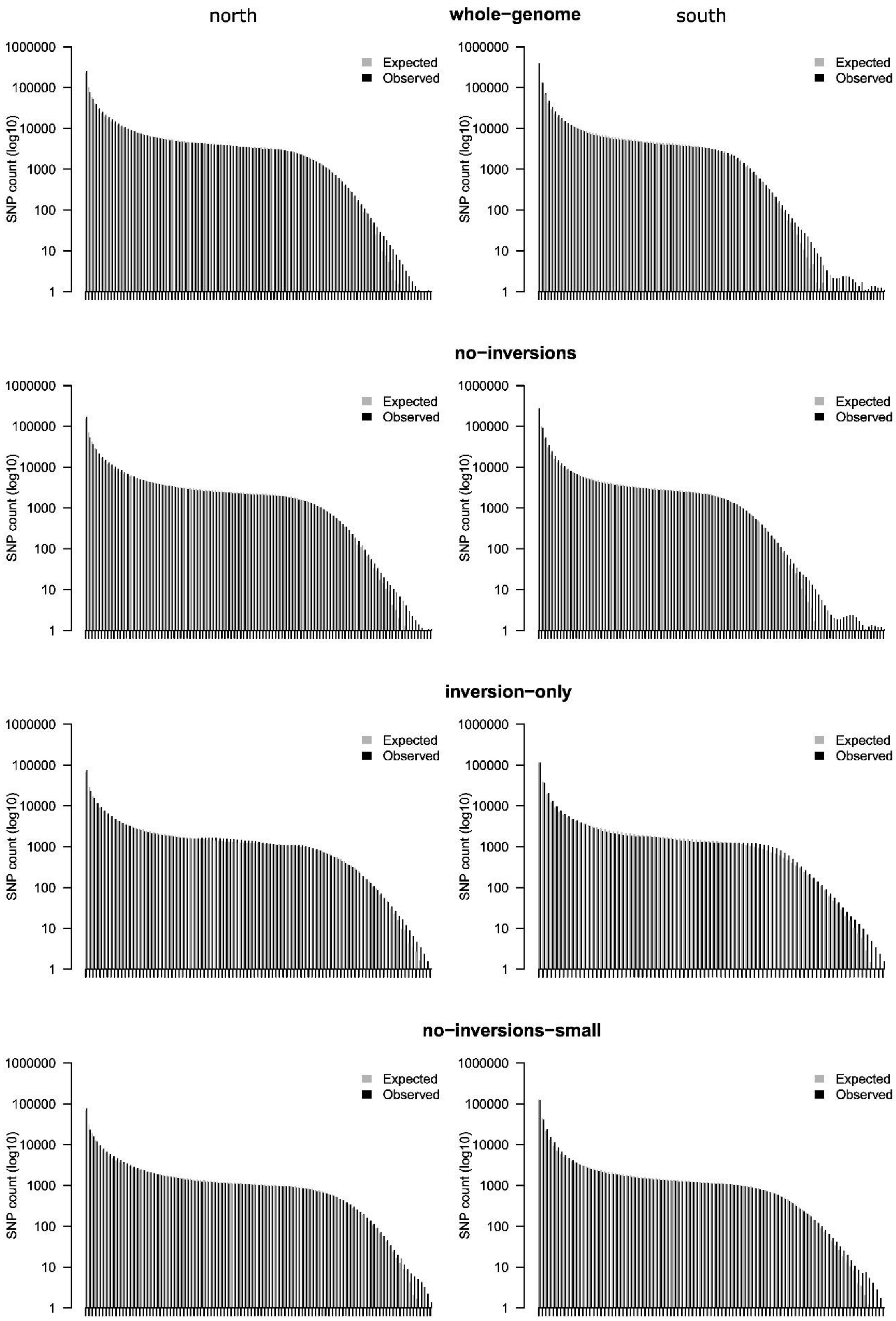
north
whole−genome
south
no−inversions
inversion−only
no−inversions−small
SNP count (log10)
1000000
100000
10000
1000
100
10
1
Expected
Observed

Figure S7. Expected one dimensional site frequency spectra (1D SFS), generated under the SC demographic model, compared with observed 1D SFS across four genomic datasets: whole-genome, no-inversions, inversions-only, and no-inversions-small. Each dataset is represented by two panels for northern (left) and southern (right) populations. Bars show SNP counts per allele frequency bin; only bins with at least one count are shown. Observed values are shown in black and expected values in grey.

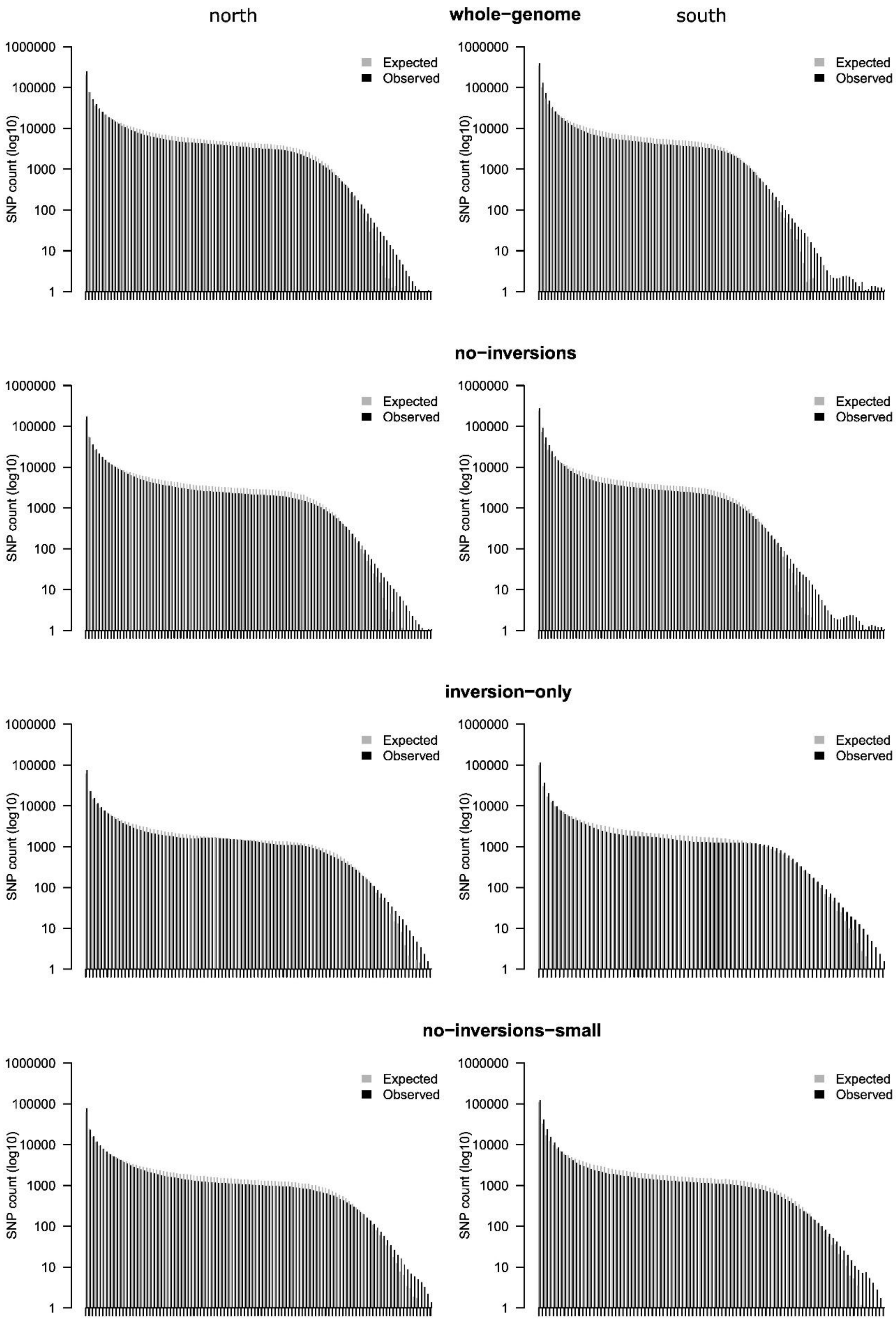
north
whole−genome
south
no−inversions
inversion−only
no−inversions−small
Expected
Observed
SNP count (log10)
1000000
100000
10000
1000
100
10
1

Figure S8. Expected one dimensional site frequency spectra (1D SFS), generated under the ISODE demographic model, compared with observed 1D SFS across four genomic datasets: whole-genome, no-inversions, inversions-only, and no-inversions-small. Each dataset is represented by two panels for northern (left) and southern (right) populations. Bars show SNP counts per allele frequency bin; only bins with at least one count are shown. Observed values are shown in black and expected values in grey.

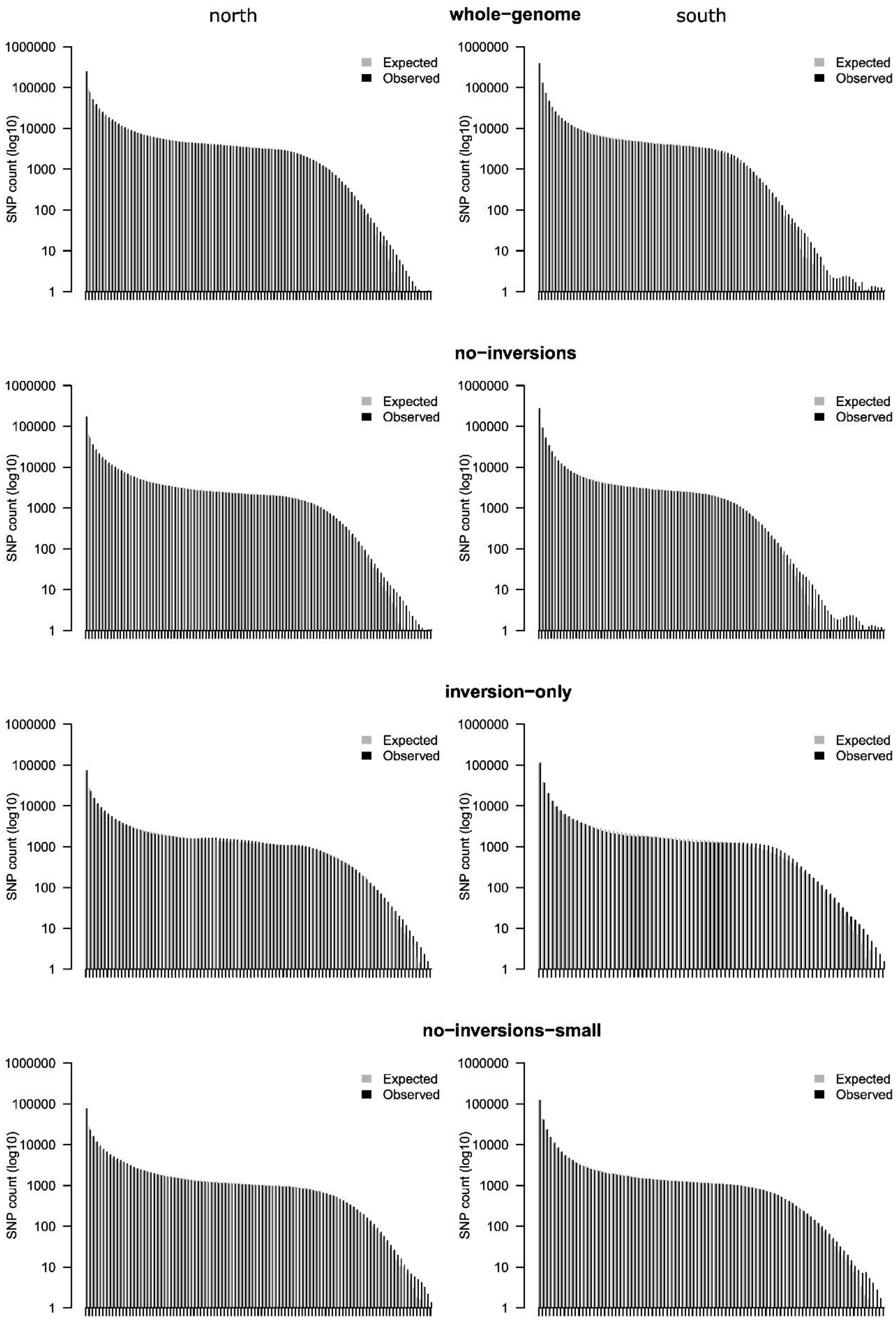

north
whole−genome
south
no−inversions
inversion−only
no−inversions−small
SNP count (log10)
1000000
100000
10000
1000
100
10
1
Expected
Observed

Figure S9. Expected one dimensional site frequency spectra (1D SFS), generated under the IMDE demographic model, compared with observed 1D SFS across four genomic datasets: whole-genome, no-inversions, inversions-only, and no-inversions-small. Each dataset is represented by two panels for northern (left) and southern (right) populations. Bars show SNP counts per allele frequency bin; only bins with at least one count are shown. Observed values are shown in black and expected values in grey.

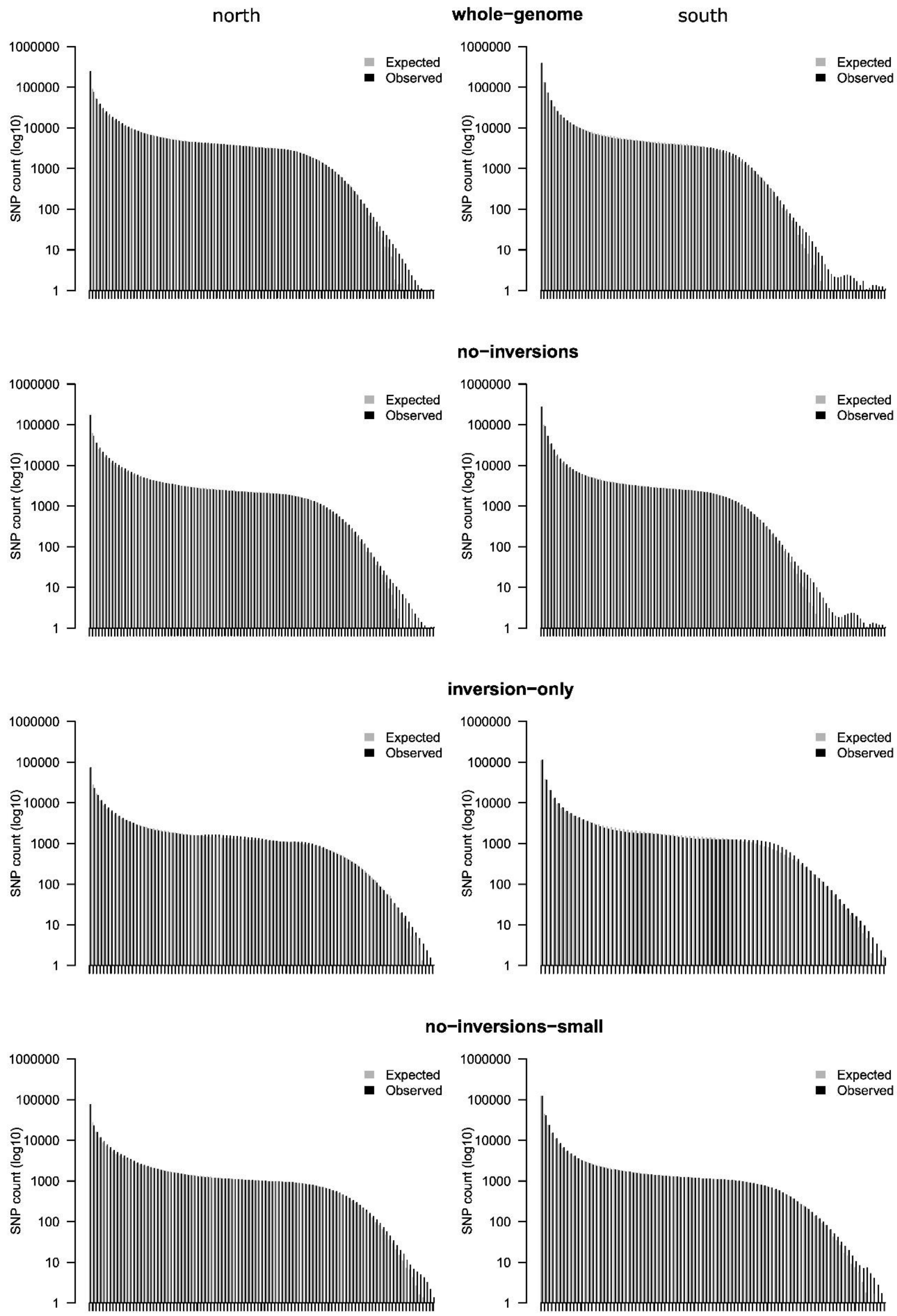
north
whole−genome
south
no−inversions
inversion−only
no−inversions−small
SNP count (log10)
1000000
100000
10000
1000
100
10
1
Expected
Observed

Figure S10. Expected one dimensional site frequency spectra (1D SFS), generated under the SCDE demographic model, compared with observed 1D SFS across four genomic datasets: whole-genome, no-inversions, inversions-only, and no-inversions-small. Each dataset is represented by two panels for northern (left) and southern (right) populations. Bars show SNP counts per allele frequency bin; only bins with at least one count are shown. Observed values are shown in black and expected values in grey.

Table S1. Sampling sites. Population ID - 3-letter population abbreviation, N - number of individuals analysed, Population name - full name of sampling site, Group - assignment to genetic group, i.e. S, C, P and N for South, Carpathian, Polish and North, respectively.

| Population ID | N | Population name | Latitude | Longitude | Country | Group |
|---|---|---|---|---|---|---|
| ABE | 13 | Abetone | 44.148 | 10.663 | Italy | South |
| KAL | 15 | Kals | 47.012 | 12.646 | Austria | South |
| LIN | 14 | Linz | 48.092 | 13.874 | Austria | South |
| BAW | 11 | Bawarian Forest | 48.960 | 13.395 | Germany | South |
| TRE | 13 | Třebíč | 49.212 | 15.879 | Czech | South |
| STE | 14 | Steigerwald | 49.622 | 10.263 | Germany | South |
| BIL | 14 | Bílkovice | 49.761 | 14.848 | Czech | South |
| SLA | 15 | Slanic | 45.250 | 25.925 | Romania | Carpathian |
| MOI | 14 | Moinesti | 46.510 | 26.449 | Romania | Carpathian |
| CAL | 15 | Calimani | 47.163 | 25.221 | Romania | Carpathian |
| ROZ | 14 | Roztocze | 50.508 | 22.786 | Poland | Poland |
| GOS | 14 | Gościnno | 54.047 | 15.657 | Poland | Poland |
| LUB | 14 | Lubaszki | 54.057 | 17.556 | Poland | Poland |
| BOR | 12 | Borki | 54.090 | 21.912 | Poland | Poland |
| TON | 13 | Tönnersjö | 56.643 | 13.070 | Sweden | North |
| ASA | 14 | Asa | 57.165 | 14.783 | Sweden | North |
| AAS | 13 | As | 59.667 | 10.793 | Norway | North |
| SIL | 14 | Siljanfors | 60.757 | 14.066 | Sweden | North |
| LAN | 13 | Länsi | 61.723 | 23.633 | Finland | North |
| EFI | 13 | Eastern Finland | 62.492 | 30.010 | Finland | North |
| STJ | 13 | Stjørdal | 63.469 | 10.918 | Norway | North |
| SVA | 13 | Svartberget | 64.236 | 19.570 | Sweden | North |
| MEL | 14 | Mellakoski | 66.399 | 24.440 | Finland | North |

Table S2. Reference genome sizes (before and after excluding genes and repeats), and the number of variants retained after quality filtering and included in the site frequency spectra, across the datasets.

| | Reference size [bp] | Reference size without genes & repeats [bp] | Number of variants after filtering | Number of variants in SFS |
|---|---|---|---|---|
| whole-genome | 162 946 098 | 44 204 527 | 1 925 226 | 1 299 931 |
| no-inversions | 118 178 221 | 31 149 941 | 1 343 318 | 910 605 |
| inversions-only | 44 767 877 | 13 054 586 | 581 908 | 391 178 |

| | | | | |
|---|---|---|---|---|
| no-inversions-small | 44 767 877 | 12 877 239 | 600 607 | 410 438 |

Table S3. Summary population genetic statistics (mean) calculated in 100 kb windows for the various datasets. Standard deviations (SD) are given in brackets. $F_{ST}$ and $d_{XY}$ were calculated between south and north genetic groups.

| | **$\pi$** | **Tajima's D** | **LD** | **$F_{ST}$** | **$d_{XY}$** |
|---|---|---|---|---|---|
| whole-genome | 0.0069 (0.0022) | -1.01 (0.357) | 0.0341 (0.0693) | 0.0223 (0.0259) | 0.0086 (0.0028) |
| no-inversions | 0.0067 (0.0024) | -1.08 (0.339) | 0.0156 (0.0478) | 0.0218 (0.0289) | 0.0085 (0.0031) |
| inversions-only | 0.0072 (0.0016) | -0.86 (0.353) | 0.0790 (0.0900) | 0.0237 (0.0167) | 0.0089 (0.0019) |
| no-inversions-small | 0.0072 (0.0014) | -1.07 (0.272) | 0.0126 (0.0301) | 0.0177 (0.0095) | 0.0091 (0.0016) |

Table S4. Pairwise $F_{ST}$ matrix calculated between all population pairs. Population abbreviations as assigned in Table S1.

| | ABE | KAL | LIN | BAW | TRE | STE | BIL | SLA | MOI | CAL | ROZ | GOS | LUB | BOR | TON | ASA | AAS | SIL | LAN | EFI | STJ | SVA | MEL |
|---|---|---|---|---|---|---|---|---|---|---|---|---|---|---|---|---|---|---|---|---|---|---|---|
| **ABE** | .000 | | | | | | | | | | | | | | | | | | | | | | |
| **KAL** | .003 | .000 | | | | | | | | | | | | | | | | | | | | | |
| **LIN** | .005 | .005 | .000 | | | | | | | | | | | | | | | | | | | | |
| **BAW** | .003 | .003 | .000 | .000 | | | | | | | | | | | | | | | | | | | |
| **TRE** | .009 | .009 | .004 | .006 | .000 | | | | | | | | | | | | | | | | | | |
| **STE** | .005 | .005 | .000 | .000 | .006 | .000 | | | | | | | | | | | | | | | | | |
| **BIL** | .014 | .013 | .007 | .011 | .001 | .011 | .000 | | | | | | | | | | | | | | | | |
| **SLA** | .010 | .012 | .006 | .006 | .013 | .003 | .020 | .000 | | | | | | | | | | | | | | | |
| **MOI** | .010 | .013 | .009 | .009 | .020 | .005 | .025 | .003 | .000 | | | | | | | | | | | | | | |
| **CAL** | .007 | .009 | .005 | .003 | .013 | .001 | .019 | .000 | .002 | .000 | | | | | | | | | | | | | |
| **ROZ** | .013 | .015 | .007 | .008 | .004 | .009 | .003 | .013 | .018 | .013 | .000 | | | | | | | | | | | | |
| **GOS** | .014 | .017 | .009 | .011 | .006 | .010 | .008 | .013 | .014 | .013 | .002 | .000 | | | | | | | | | | | |
| **LUB** | .021 | .021 | .015 | .015 | .012 | .015 | .011 | .020 | .022 | .019 | .003 | -.001 | .000 | | | | | | | | | | |
| **BOR** | .021 | .023 | .016 | .014 | .014 | .016 | .018 | .020 | .025 | .018 | .007 | .005 | .003 | .000 | | | | | | | | | |
| **TON** | .027 | .028 | .021 | .023 | .020 | .019 | .023 | .020 | .022 | .020 | .011 | .005 | .004 | .005 | .000 | | | | | | | | |
| **ASA** | .030 | .034 | .023 | .026 | .022 | .023 | .023 | .026 | .028 | .025 | .012 | .006 | .005 | .006 | .003 | .000 | | | | | | | |
| **AAS** | .029 | .029 | .024 | .025 | .020 | .023 | .022 | .026 | .028 | .025 | .012 | .007 | .004 | .004 | .000 | .002 | .000 | | | | | | |
| **SIL** | .029 | .031 | .025 | .024 | .021 | .024 | .021 | .029 | .029 | .026 | .011 | .006 | .002 | .003 | .001 | .000 | -.001 | .000 | | | | | |
| **LAN** | .026 | .026 | .024 | .023 | .022 | .021 | .025 | .024 | .021 | .021 | .014 | .006 | .004 | .005 | .001 | .004 | .001 | .001 | .000 | | | | |
| **EFI** | .028 | .032 | .024 | .025 | .022 | .022 | .025 | .024 | .024 | .023 | .013 | .003 | .004 | .005 | .001 | .002 | .003 | .001 | -.001 | .000 | | | |
| **STJ** | .028 | .030 | .024 | .025 | .025 | .021 | .028 | .024 | .022 | .022 | .015 | .007 | .006 | .008 | -.001 | .003 | .001 | .001 | .000 | .001 | .000 | | |
| **SVA** | .030 | .030 | .028 | .026 | .031 | .025 | .033 | .026 | .021 | .024 | .019 | .010 | .010 | .012 | .002 | .008 | .006 | .005 | .001 | .003 | .000 | .000 | |
| **MEL** | .024 | .026 | .020 | .019 | .023 | .018 | .028 | .020 | .019 | .017 | .014 | .008 | .007 | .003 | .001 | .004 | .003 | .004 | -.001 | .001 | .000 | .002 | .000 |

Table S5. Deviations from Hardy-Weinberg equilibrium tested per inversion. Undivided refers to all individuals used in PSMC (see Materials and methods). Higher, Similar and Lower refer to groups of individuals expressing particular deviations from the Ne estimate obtained for no-inversions-small dataset. p - uncorrected p-value from Fisher exact test. FDR - p-value corrected for multiple testing. Inversion names as assigned in Mykhailenko et al. 2024.

| **Inversion** | **Undivided** | | **Higher** | | **Similar** | | **Lower** | |
|---|---|---|---|---|---|---|---|---|
| | p | FDR | p | FDR | p | FDR | p | FDR |
| Inv2 | 0.056 | 0.389 | 0.082 | 0.357 | 1.000 | 1.000 | NA | NA |
| Inv3 | 0.166 | 0.582 | 0.184 | 0.597 | 0.676 | 1.000 | NA | NA |
| Inv5 | 0.276 | 0.734 | 1.000 | 1.000 | 0.654 | 1.000 | 0.167 | 0.909 |
| Inv6 | 0.136 | 0.542 | **0.049** | 0.255 | 1.000 | 1.000 | 1.000 | 1.000 |
| Inv7.1 | 0.370 | 0.759 | 0.407 | 0.781 | 0.799 | 1.000 | 1.000 | 1.000 |
| Inv7.2 | 0.745 | 1.000 | 1.000 | 1.000 | 0.416 | 1.000 | 1.000 | 1.000 |

| | | | | | | | | |
|---|---|---|---|---|---|---|---|---|
| Inv10 | 0.380 | 0.759 | 0.101 | 0.373 | 0.803 | 1.000 | 1.000 | 1.000 |
| Inv12 | 0.104 | 0.485 | 0.489 | 0.848 | 0.135 | 0.756 | NA | NA |
| Inv13 | 0.242 | 0.734 | 0.303 | 0.781 | 1.000 | 1.000 | 1.000 | 1.000 |
| Inv14.1 | 0.722 | 1.000 | 0.596 | 0.911 | 0.795 | 1.000 | 0.594 | 1.000 |
| Inv14.2 | 0.315 | 0.734 | 0.702 | 1.000 | 0.250 | 0.932 | NA | NA |
| Inv14.3 | 0.290 | 0.734 | 0.417 | 0.781 | 0.317 | 0.986 | 1.000 | 1.000 |
| Inv14.4 | 1.000 | 1.000 | 1.000 | 1.000 | 1.000 | 1.000 | 1.000 | 1.000 |
| Inv14.5 | 0.856 | 1.000 | 0.399 | 0.781 | 0.791 | 1.000 | 1.000 | 1.000 |
| Inv14.6 | 1.000 | 1.000 | 1.000 | 1.000 | 1.000 | 1.000 | 1.000 | 1.000 |
| Inv15 | **0.024** | 0.361 | **0.007** | 0.057 | 0.533 | 1.000 | 1.000 | 1.000 |
| Inv16.1 | 0.698 | 1.000 | **0.005** | 0.057 | **0.038** | 0.269 | 0.289 | 0.909 |
| Inv16.2 | **0.039** | 0.361 | NA | NA | **0.024** | 0.269 | NA | NA |
| Inv17 | 1.000 | 1.000 | 0.421 | 0.781 | 0.604 | 1.000 | 0.282 | 0.909 |
| Inv18 | 0.956 | 1.000 | **0.044** | 0.255 | 0.669 | 1.000 | 1.000 | 1.000 |
| Inv22.1 | 0.710 | 1.000 | 0.216 | 0.623 | 0.796 | 1.000 | 1.000 | 1.000 |
| Inv22.2 | 0.851 | 1.000 | 1.000 | 1.000 | 0.266 | 0.932 | 0.548 | 1.000 |
| Inv22.3 | 0.072 | 0.402 | 0.842 | 1.000 | 0.201 | 0.932 | 0.225 | 0.909 |
| Inv22.4 | 0.483 | 0.902 | 0.756 | 1.000 | 0.650 | 1.000 | 0.289 | 0.909 |
| Inv22.5 | 1.000 | 1.000 | 1.000 | 1.000 | 1.000 | 1.000 | 1.000 | 1.000 |
| Inv23.1 | 0.698 | 1.000 | **0.005** | 0.057 | **0.038** | 0.269 | 0.289 | 0.909 |
| Inv23.2 | **0.039** | 0.361 | NA | NA | **0.024** | 0.269 | NA | NA |
| Inv26 | 0.860 | 1.000 | 0.589 | 0.911 | 0.445 | 1.000 | 0.205 | 0.909 |
